# Movement characteristics impact decision-making and vice versa

**DOI:** 10.1101/2022.02.02.478832

**Authors:** Thomas Carsten, Fanny Fievez, Julie Duque

**Author notes:** Corresponding author: Thomas Carsten. **Author contributions** Thomas Carsten: Conceptualization, Methodology, Software, Formal Analysis, Investigation, Data Curation, Writing – Original Draft Preparation, Visualization, Funding Acquisition Fanny Fievez: Conceptualization, Methodology, Writing – Review & Editing Julie Duque: Conceptualization, Methodology, Writing – Review & Editing, Supervision, Funding Acquisition. **Data availability** All data and accompanying scripts will be deposited on the Open Science framework for free access before peer-reviewed publication of this manuscript: https://osf.io/dk7a2/.

## Abstract

Previous studies suggest that humans are capable of coregulating the speed of decisions and movements if promoted by task incentives. It is unclear however whether such behavior is inherent to the process of translating decisional information into movements, beyond posing a valid strategy in some task contexts. Therefore, in a behavioral online study we imposed time constraints to either decision or movement -phases of a sensorimotor task, ensuring that coregulating decisions and movements was not promoted by task incentives. We found that participants indeed moved faster when fast decisions were promoted and decided faster when subsequent movements had to be executed swiftly. Furthermore, inflicting faster movements seems to alter decision-making in a similar fashion as conditions promoting faster decisions: In both fast-decision and fast-movement blocks, decisions relied more strongly on information presented shortly rather than long before movement onset. Taken together, these findings suggest that decisions not only impact movement characteristics, but that properties of movement impact the time and manner with which decisions are made. We interpret these behavioral results in the context of *embodied decision-making*, whereby shared neural mechanisms may not only enable faster movements but also assist in making decisions in less time.

## Introduction

When animals hunt, they may have to decide quickly whether turning left or right allows cutting into their prey’s path, before putting this plan into action. Traditional views have understood this sensorimotor process as consisting of at least two functionally distinct brain processes (e.g. Ratcliff & McKoon, 2008; Sternberg, 1969): ‘Deciding’ regards the comparison and selection between expected outcomes based on sensory information, whereas ‘moving’ reflects the interaction with the environment to achieve the favored outcome. These two processes have been associated with different brain structures (Farrar et al., 2018 Welniarz et al., 2022) and different neurophysiological signatures (Desender et al., 2021; Kelly & O’Connell, 2015; Wessel, 2020). Perhaps as a result of this predominant view, research on decision-making and movement specification have been covered by different scientific ‘disciplines’ (Cisek & Pastor-Bernier, 2014), leaving a blind spot on the interplay of both aspects of sensorimotor processing.

However, over the course of the past ten years or so, increased interest in the interaction between decisions and movements has generated findings which challenge the view that these processes operate independently. For instance, properties of decisions such as reward expectancy (Klein-Flügge & Bestmann, 2012; Shadmehr et al., 2019), choice preference (Calderon et al., 2018; de Lange et al., 2013; Klein et al., 2012), conflict anticipation (Derosiere et al., 2018; Duque et al., 2016) and time pressure (Derosiere et al., 2021; Kelly et al., 2021; Murphy et al., 2016; Steinemann et al., 2018) systematically change activity in motor areas of the brain, and alter kinematics of movements expressing those decisions (Freeman, 2018; Shadmehr et al., 2019; Spieser et al., 2017; Thura, 2020). Likewise, kinematic properties of movements such as required time (Burk et al., 2014; Reynaud et al., 2020) and energetic costs (Hagura et al., 2017; Marcos et al., 2015; Morel et al., 2017) are factored into decisions. Taken together, these findings challenge the view that decisions and movements pose fully functionally distinct aspects of sensorimotor processing (Barsalou, 2008).

Such interplay may reflect adjustments concerning both decisions and movements in order to increase the net payoff of behavior. For example, biomechanical costs of movement may be weighed against its expected outcome in order to justify its effort expenditure (Cos et al., 2014; Morel et al., 2017; Shadmehr et al., 2016). Similarly, under time pressure, reducing both the time to decide and to move shortens the overall time required to respond (Spieser et al., 2017). Taken together, animals, including humans, likely aim to maximize ‘capture rate’ (Reynaud et al., 2020; Shadmehr et al., 2019; Thura, 2020; Yoon et al., 2018) by minimizing time and effort required to obtain reward, leading to adaptations concerning both decisions and movements.

If animal behavior in terms of decisions and movements serves a common function to secure capture rate and hence survival, then a close consensus between decisions and movements may be naturally promoted by the organization of the sensorimotor system (Cisek, 2019). In other words, decisions and movements may not be fully functionally distinct brain processes, as they serve a common function and hence likely rely on shared brain circuity through evolution. As such, movements may be considered *expressed* or *embodied* decisions in that they establish preferred outcomes through interaction with the environment. From this theoretical perspective, termed *embodied decision-making*, a gradual shift in functionality exists between anatomically distinct brain areas, transforming abstract- decisional considerations of future consequences to their concrete implementation through movement (Cisek & Pastor-Bernier, 2014; Lepora & Pezzulo, 2015; Wispinski et al., 2020; Yoo & Hayden, 2018). The key difference to more traditional viewpoints is the acknowledgement of reciprocal, continuous and interactive communication between brain areas associated with decisional and motor processes (Wispinski et al., 2020), leading to a distributed consensus about the preferred course of action, reflecting both abstract decisional considerations of future consequences and concrete motor requirements for the implementation of the response (Yoo & Hayden, 2018). As such, embodied decision-making accounts predict a close functional relationship between decisions and movements, whereby movement kinematics depend on decisional information and the integration of information into decisions is fundamentally constrained by movement requirements.

It was the goal of this study to seek behavioral evidence for embodied decision-making, which has remained largely theoretical despite of its appeal. Although previous studies in this regard largely demonstrated how decisions and (preparation of) movements change under time pressure (Jaśkowski et al., 2000; Kelly et al., 2021; Murphy et al., 2016; Pastötter et al., 2012; Spieser et al., 2017, 2018; Steinemann et al., 2018; Thura, 2020), the nature of these tasks actively promoted a joint reduction of time taken to decide and move, to reduce overall response times. However, from an embodied decision-making perspective a mutual dependence of decisions and movements may reflect an inherent property of the sensorimotor system. It should therefore be observable even if there is no task incentive linking decisions and movements. Likewise, it implies that such interdependence sustains even when time pressure is limited towards *either* decisions or movements, rather than towards both at the same time as in previous studies.

With a behavioral online study run on 62 participants, we tested four predictions grounded in the *embodied decision-making* framework. As a first hypothesis, people should tend to move quicker if they have less time to decide, even if such behavior is not required, nor advantageous. As a second hypothesis, conversely, people should tend to decide faster as they need to perform faster movements, again even if not profitable. Third and fourth predictions were derived from the idea that embodied decisions emerge from a distributed consensus between decisional and movement concerns (Cisek, 2012; Yoo & Hayden, 2018). Such consensus implies that the preparation to move may be seen as gradual commitment towards a decision, as readiness to initiate movement may increase costs of revoking movement and hence changing the decision (Lepora & Pezzulo, 2015). Therefore, it was our third prediction that decisions should rely more on earlier information as compared to information presented shortly before movement onset. This is conceptually similar to findings showing that committing to a decision reduces sensitivity towards subsequent information (Bronfman et al., 2015; Talluri et al., 2021), however preparation of movement itself should reflect such gradual commitment. As a final fourth prediction, such *primacy effect* with which novel information is integrated into decisions should be strongest when high motor control is required, such as when performing fast as compared to slow tapping (Jäncke, Peters, et al., 1998; Jäncke, Specht, et al., 1998; Lutz et al., 2004). This is because costs of revoking movement and hence changing the decision should increase under high motor control (Burk et al., 2014), leading to less consideration of later information for embodied decisions. Especially this fourth hypothesis allows a strong test for embodied decision-making, as it predicts that people arrive differently at their decisions based on the movement required to express those.

In summary, we sought evidence for embodied decision-making by testing that a) the time taken to decide and move constrain each other even if not adaptive in the context of the task and b) integration of information into decisions over time is constrained by the movement required to express those decisions. We found such evidence by imposing time constraints either to decision phases or movement phases of a sensorimotor task. Participants indeed reduced both the time to decide and to move although time constraints regarded only either decisions or movements. Remarkably, most participants who displayed larger instructed time savings in either decision or movement duration also speeded up corresponding movements or decisions more, respectively, although not profitable in this task. Furthermore, the influence of novel information on ongoing decisions fundamentally depended on the timing relative to the motor response: Contrary to our expectations and likely driven by task design, novel information presented shortly before movement initiation was taken into account more than earlier information. This *recency effect* was however accentuated when participants performed faster movements, providing strong evidence that movement characteristics affect decision-making. Taken together, these findings support the idea that decisions and movements are not fully functionally distinct brain processes, but condition each other even when not required in the context of the task.

## Results

Sixty-two participants performed two separate sessions of a behavioral online experiment from home. We administered an adaptation of the Tokens task, which has been widely used to investigate the effect of time pressure on sensorimotor processes in humans (Cisek et al., 2009; Derosiere et al., 2019; Reynaud et al., 2020; Lunazzi et al., 2021; Thura, 2020). Each trial consisted of a dedicated decision and movement phase in order to measure the time participants would allocate to deciding and moving (see Figure 1). In the decision phase, participants decided which one of two bananas would outgrow the other; both grew at a non-constant speed over time. Their decision had to be indicated by a left- or rightward key press, which initiated the subsequent movement phase. Here, participants continued tapping this key to move a caterpillar upwards. The caterpillar moved faster with faster key tapping. This basic task template was modified in a ‘decision session’ to encourage fast or slow decisions in separate blocks, with no constraints on tapping speed. In a separate ‘movement session’, fast or slow finger tapping was required, with no restrictions on decision speed. Comparing finger tapping speed in the movement phase between blocks of fast and slow decisions and comparing decision duration in the decision phase between blocks of fast and slow tapping allowed to test the hypothesis that the duration of decisions and movements impact each other, even if not profitable in the context of the task.

**Figure 1.**
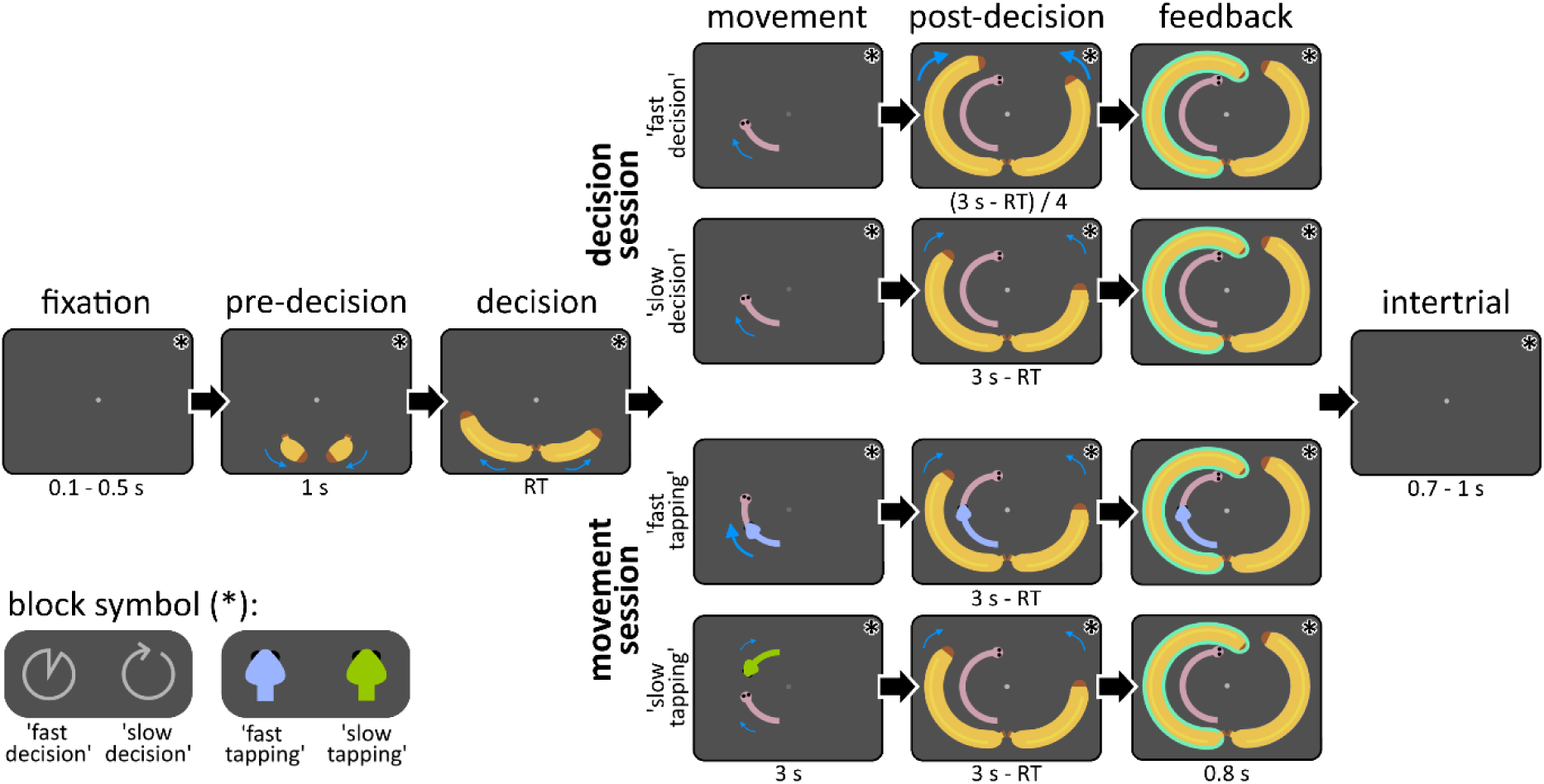
Trial structure. After some time of fixation, baby bananas moved symmetrically into position below the fixation circle, telegraphing the onset of the decision phase. The latter was characterized by asymmetrical growth of bananas, during which participants had to indicate which banana would outgrow the other with a key press. After this decision, the same key had to be pressed four more times in the movement phase to move a caterpillar into the corresponding direction. Here, the required tapping speed depended on the experimental condition: Fast-tapping blocks required participants to tap fast to avoid a chasing snake, whereas slow-tapping blocks required slow tapping to avoid a retreating snake. In contrast, fast-decision and slow-decision conditions had no specific finger tapping speed requirements but required to put the emphasis on either decision speed or accuracy, respectively. In the post-decision phase, bananas finished their growth trajectories until a full circle was covered. Here, bananas grew at same speed as in the decision phase, with the exception of the fast-decision condition, during which bananas grew four times faster. Afterwards, decision feedback was given by highlighting the chosen banana either in green (correct) or red (incorrect). A final intertrial interval preceded the next trial. Blue arrows depict movement trajectories of the stimuli, with thicker and longer arrows reflecting faster movement. Blue arrows are shown for illustrative purposes and were not visible to participants. Symbols were shown in the upper right corner of the screen (indicated by asterisks), as constant reminder of one of the four experimental block conditions. ‘s’ = seconds, ‘RT’ = reaction time.

### When participants are required to decide faster, they also move faster

Participants followed the instructions consistently and adjusted their decision duration in fast and slow decision-blocks. As seen in Figure 2A, participants took about 672 ms (*95%- Confidence Interval [95%-CI]*: 585-733 ms) less to decide which banana to choose in fast- decision (*median* = 1057 ms, *Median Absolute Deviation [MAD]* = 385 ms) as compared to slow-decision blocks (*median* = 1767 ms, *MAD* = 345 ms, *S* = 2, *p* < 10^-13^). This came at the price of reducing probability of making the correct decision by 16% (*95%-CI*: 13-19%) from a median of 84% (*MAD* = 6%) in slow-decision to 66% (*MAD* = 8%) in fast-decision blocks (*S* = 4, *p* < 10^-11^), reflecting a shift in speed-accuracy tradeoff (see Supplemental Figure S3A). As a consequence, there was a reduction in the proportion of correct decisions by 14% (*95%-CI*: 10-19%) from slow-decision (*median* = 91%, *MAD* = 3%) to fast-decision blocks (*median* = 76%, *MAD* = 13%, *S* = 5, *p* < 10^-10^, see Supplemental Figure S3B).

**Figure 2.**
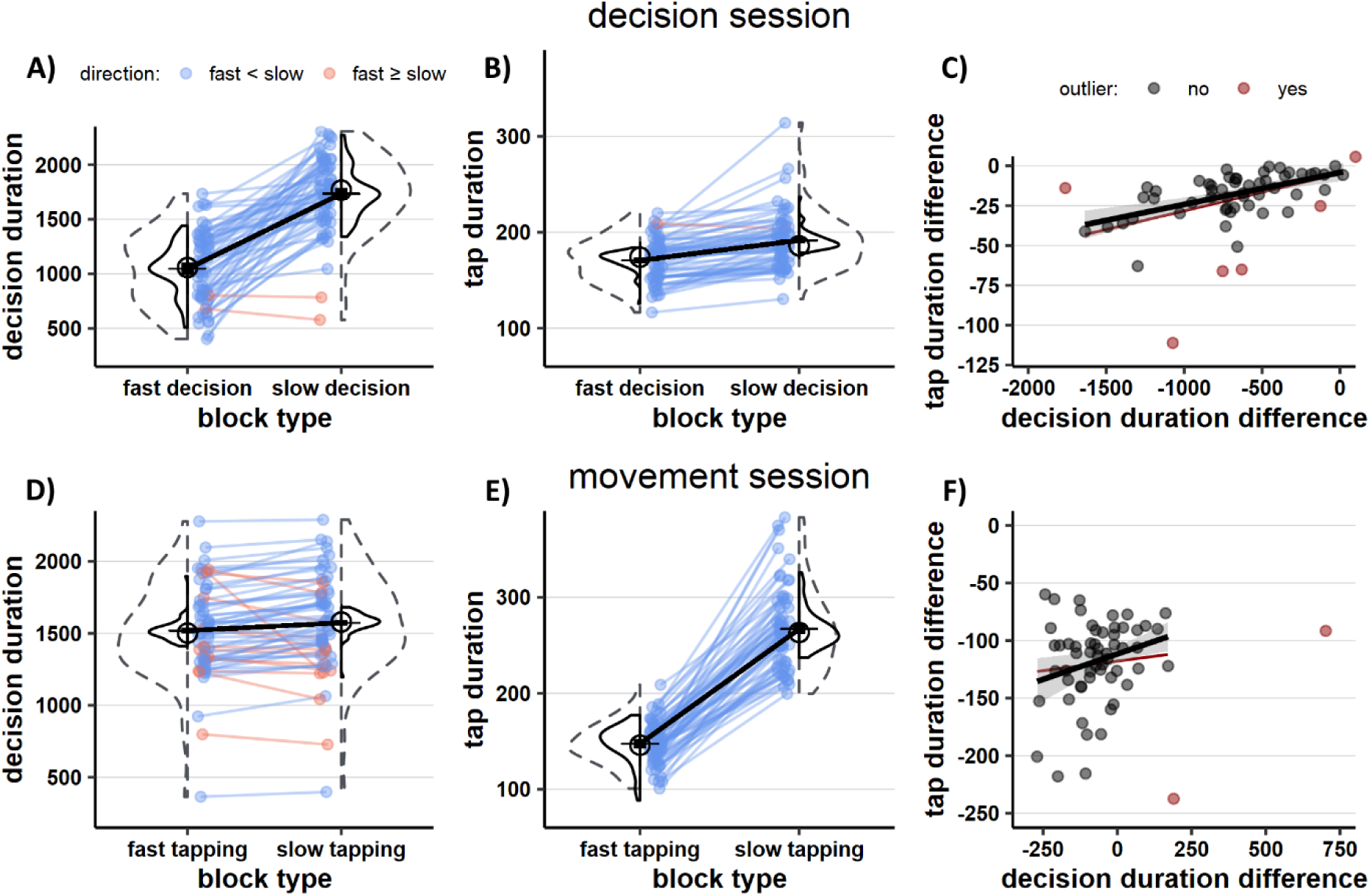
Behavioral results. The upper graphs (A-C) depict data for the decision session; three lower graphs (D-F) are for the movement session. **A)** As instructed, participants changed their decision duration between blocks in the decision session. **B)** Although not profitable in the context of the task, participants tapped faster in fast-decision blocks as compared to slow-decision blocks. **C)** There was a linear relationship between the instructed change in decision duration and the resulting change in tap duration across participants. This linear relationship was significant both when considering the entire sample (red line), as well as the sample after removing outliers (red data points and black line, gray area reflects 95%- CI). **D)** Although not profitable in the context of the task, participants decided faster in fast-tapping blocks as compared to slow-tapping blocks. **E)** As instructed, participants changed their tap duration between blocks in the movement session. **F)** There was no linear relationship between the instructed change in tap duration and the resulting change in decision duration across participants when considering the whole sample (red line). However, there was a significant relationship after removing outliers (red data points and black line). **A, B, D, E)** Individual data is shown as colored points connected by a line, with blue data indicating individuals with numerically lower values in fast as compared to slow decision or movement blocks and orange indicating the opposite. Dashed histograms indicate the distribution of data points for each condition. Solid histograms reflect the same distributions after centering subjects around the group mean to remove between-subjects variance unrelated to the experimental manipulation (Cousineau, 2005). The average effect on the group level is shown by plotting the mean (black horizontal lines), connected by a black line. Black bars around the mean reflect its within-subject 95% confidence interval (Cousineau, 2005). The group median of each condition is shown as circle.

As predicted, this reduction in decision duration came with increased tapping speed in the movement phase following the decision (see Figure 2B). The average duration of the four individual key taps was reduced by circa 16 ms (*95%-CI*: 12-20 ms) from slow-decision (*median* = 186 ms, *MAD* = 27 ms) to fast-decision blocks (*median* = 174 ms, *MAD* = 26 ms, *S* = 1, *p* < 10^-15^). This experimental effect was significant for each of the four finger taps in the movement phase (all *p*’s < 10^-9^, see Supplemental Figure S4A). Post hoc comparisons did not indicate that this experimental effect was larger for earlier finger taps (all *p*’s ≥ .44), although there was a significant BLOCK TYPE x TAP NUMBER interaction (*F*(2.29, 130.80) = 10.15, *p* < 10^-4^ and η^2^_p_= .15, see Figure S4A) in a repeated-measures ANOVA. Similar results were obtained for this ANOVA after removing extreme values lying three median absolute deviations (*MAD*s) outside the group median to test for robustness of these findings. Remarkably, Figure 2C shows that those participants with the largest reduction in decision duration were also those who increased tapping speed the most between slow and fast decision-blocks (*t* = 4.34, *p* < 10^-4^ and *r* = .50). A similar linear relationship was observed after removing six influential values from this analysis (i.e. those with a Cook’s Distance three times larger than the mean, red dots in Figure 2C; *t* = 5.07, *p* < 10^-5^ and *r* = .58). Virtually all finger taps in the movement phase of fast-decision (*median* = 100%, *MAD* = 0%) and slow-decision (*median* = 100%, *MAD* = 0%) blocks were performed correctly, with no difference between blocks (*S* = 6, *p* > .99, see Supplemental Figure S3C).

The interpretation that participants tapped faster when fast decisions were required was substantiated by fitting behavior to drift diffusion models. These models assume that reaction times reflect the duration of the sensorimotor process covering sensation, decision-making and response execution. By fitting different candidate models to behavior, it can be determined which aspects of this sensorimotor process most likely differed between experimental conditions. A model, which allowed the decision threshold, non-decision time and drift rate to differ between experimental conditions, described behavior best (see Table S2). The decision threshold (as proxy for changes in decision duration due to shifts in speed-accuracy tradeoff) was significantly lower in fast-decision blocks than in slow-decision blocks (Probability > 99.975%, i.e. exceeding 3999 out of 4000 posterior samples). Critically, the non-decision time (as proxy for sensorimotor delays related to initiating and executing the first key press) was also significantly reduced (Probability > 99.975%, see Table 1). Although not expected, the drift rate (as proxy for the efficiency with which a correct decision can be made) was also lowered (Probability > 99.975%), indicating that choosing under time pressure let to worse decision performance beyond deliberate shifts in speed-accuracy tradeoff (cf. Rae et al., 2014).

**Table 1.** Estimated drift diffusion model parameters per experimental condition.

| | decision threshold ($a$) | | non-decision time ($t$) | | drift rate ($v$) | |
| --- | --- | --- | --- | --- | --- | --- |
|  | mean | CI | mean | CI | mean | CI |
| slow decision (intercept) | 3.17 | 3.02<br>3.34 | 0.73 | 0.65<br>0.82 | 0.97 | 0.90<br>1.04 |
| fast decision (difference) | -1.04 | -1.22<br>-0.85 | -0.35 | -0.44<br>-0.25 | -0.42 | -0.49<br>-0.34 |
| slow tapping (intercept) | 2.87 | 2.73<br>3.01 | 0.80 | 0.72<br>0.89 | 0.91 | 0.85<br>0.98 |
| fast tapping (difference) | -0.11 | -0.21<br>-0.01 | -0.05 | -0.11<br>0.01 |  |  |

*Note.* Parameter values are based on group posteriors of the best-fitting drift diffusion models of each session. Credible intervals (CI) cover 2.5% and 97.5% percentiles of the respective posterior distribution. To estimate within-subject effects, an intercept was fitted to the slow condition and the difference thereof was fitted to the fast condition of the respective session. For the movement session, only one drift rate was fitted for both experimental conditions, as this provided better model fit than the full model (see Table S2).

### When participants are required to move faster, they also decide faster

As shown in Figure 2E, participants also followed the instructions consistently in the movement session. Average duration of each of the four key taps in the movement phase was circa 111 ms faster (*95%-CI*: 104-125 ms) when fast tapping (*median* = 146 ms, *MAD* = 22 ms) was required as compared to slow tapping (*median* = 263 ms, *MAD* = 45 ms, *S* = 0, *p* < 10^-17^). Each single finger tap was consistently faster when fast tapping was required as compared to slow tapping (all *p* < 10^-16^). However the first finger tap of the movement phase was speeded up the most from slow to fast tapping-blocks (all *p* < 10^-7^ as compared to subsequent taps), as well as the second tap being sped up more than the last (*p* = .04, with no other significant comparisons), as indicated by a BLOCK TYPE and TAP NUMBER interaction (*F*(1.22, 70.93) = 36.71, *p* < 10^-8^ and η^2^_p_ = .39, see Supplemental Figure S4B). Similar results were obtained after removing extreme values to test for robustness. Hence, the experimental manipulation was effective in shifting tapping speed of participants between blocks. The proportion of trials in which participants performed tapping movements correctly was high, however it was significantly lower in fast-tapping (*median* = 93%, *MAD* = 4%) as compared to slow-tapping blocks (*median* = 100%, *MAD* = 0%, *S* = 1, *p* < 10^-15^, see Figure S3F), consistent with the higher motor control requirement associated with the former condition.

We then tested the hypothesis that decision duration changes with tapping speed. As predicted and shown in Figure 2D, decisions were circa 73 ms (*95%-CI*: 35-98 ms) faster in fast-tapping blocks (*median* = 1502 ms, *MAD* = 296 ms) as compared to slow-tapping blocks (*median* = 1579 ms, *MAD* = 296 ms, *S* = 13, *p* < 10^-4^). The probability of making a correct decision was however not significantly different between slow-tapping blocks (*median* = 80%, *MAD* = 7%) and fast-tapping blocks (*median* = 79%, *MAD* = 7%, *S* = 24, *p =* .19, see Supplemental Figure S3D). Likewise, this did not result in a change in the proportion of correct decisions from slow-tapping (*median* = 90%, *MAD*= 4%) to fast-tapping blocks (*median* = 90%, *MAD* = 4%, *S* = 26, *p* = .43, see Supplemental Figure S3E). Across all participants, we did not find that those reducing decision duration the most, where also those who speeded up tapping the most between blocks (see Figure 2F; *t* = 1.78, *p* = .08 and *r* = .23). However, such linear relationship was present after removing two influential values based on Cook’s Distance (red dots in Figure 2F; *t* = 2.69, *p* < .01 and *r* = .34). This suggests that faster tapping is preceded by proportionally faster decisions in most, but not all participants.

Drift diffusion modelling corroborated the interpretation that requirements to tap fast shortened decisions: That is, out of five candidate models, the hypothesized model fit behavioral data best. This model allowed the decision threshold and the non-decision time to differ between experimental conditions (see Table S2). The decision threshold was significantly reduced in fast-tapping blocks (Probability = 98.30%), supporting the interpretation that movement requirements reduced the time taken to decide. The reduction in non-decision time did not reach conventional levels of significance (Probability = 95.60%), potentially since experimental constraints on tapping speed regarded the movement phase, but not the first tap during the decision phase (visualized as chasing or retreating snakes, see Figure 1), which was modelled.

### Experimental conditions influence how people arrive at their decisions over time

In line with embodied decision-making, we expected gradual commitment to a decision as corresponding movement is progressively prepared (Yoo & Hayden, 2018). Therefore, decisions should rely more on earlier evidence than evidence presented shortly before the movement. To test this, we computed temporal weighting profiles (see Figure 3). These show in how far participants’ decisions can be predicted by decisional information, depending on when in time this information is shown. Negative slopes of these profiles suggest that participants prioritized earlier over later information, i.e. a primacy effect. A positive slope suggests a recency effect, i.e. the opposite. In contrast to the hypothesized primacy effect, we found a significant recency effect in all experimental conditions (all *p*’s < 10^-15^). Indeed, visual inspection of Figure 3 suggests that decisional information had the strongest influence on the participants’ decisions when presented between 400 and 100 ms before the estimated onset of the first finger tap.

**Figure 3.**
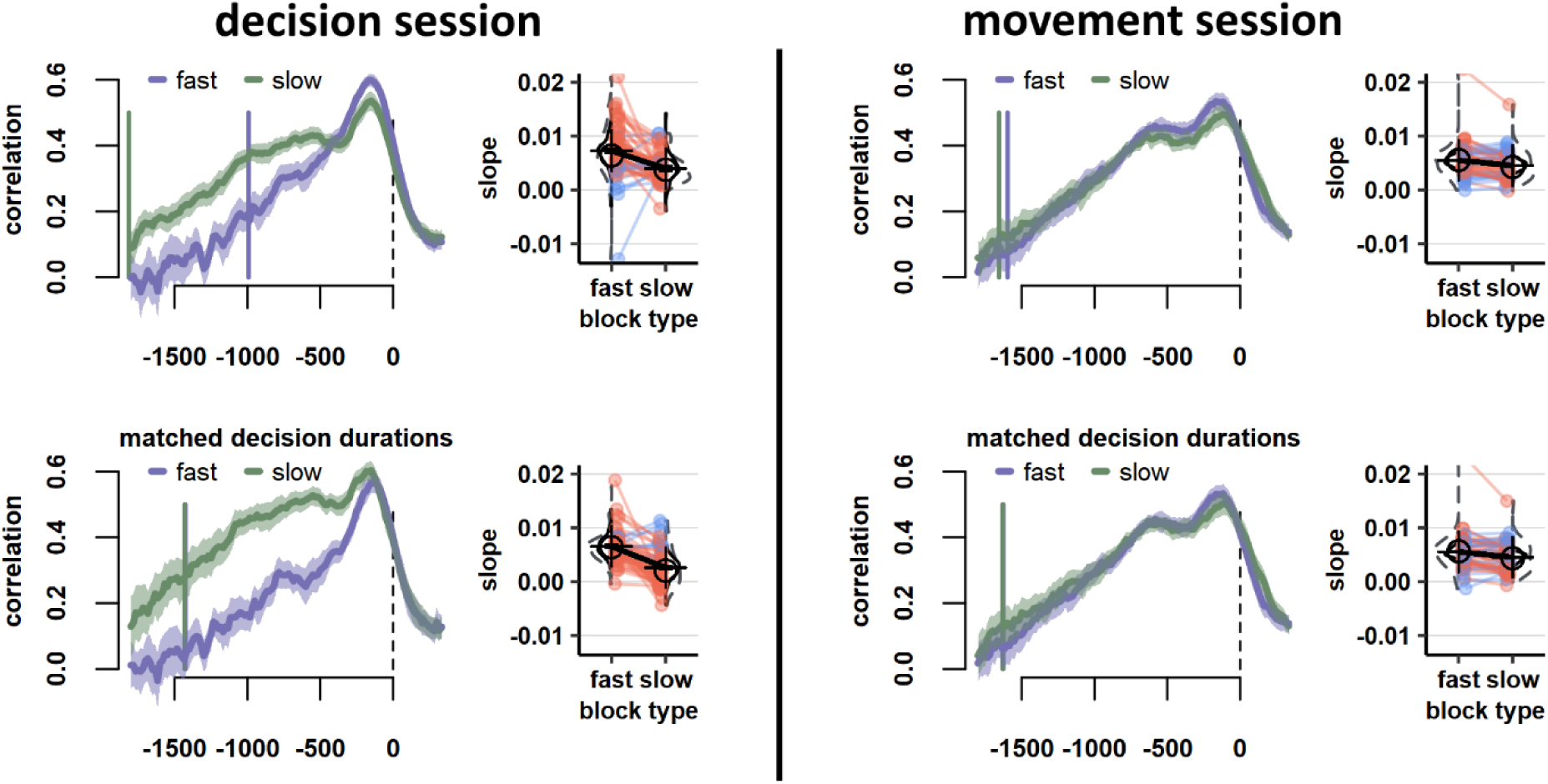
Temporal weighting profiles. Participants prioritized recent over earlier evidence when making their decisions (i.e. a recency effect), and more so when either fast decisions or fast tapping was required. Left and right columns show data from the decision session and movement session, respectively. The upper row shows the main results and the lower row shows a control analysis with matched decision durations between experimental conditions of the same session. Respective left larger line graphs show how the correlation between momentary evidence and participants’ decisions (y-axis) changes as a function of the time (x-axis, in milliseconds) relative to the estimated onset of the first finger tap (dashed vertical line, i.e. ‘response-locked’). Green lines reflect temporal weighting profiles in either slow-decision blocks (left column) or slow-tapping blocks (right column), whereas purple lines reflect either fast-decision blocks (left column) or fast-tapping blocks (right column). Line graphs depict grand averages across participants, with lighter colors depicting 95%- confidence intervals. Vertical colored solid lines represent the expected onset of the decision (phase) per experimental condition, computed as negative average median decision duration across participants. In the lower row, these vertical lines are overlapping, since decision durations were matched between experimental conditions (see Materials and Methods). Statistical tests were conducted on the basis of individual slopes of temporal weighting profiles, which are shown as smaller graphs on the respective right side. Individual slopes were extracted from the interval of negative median decision duration (estimated decision onset) to 0 (estimated decision offset). For individual slopes, red lines depict participants with a numerically stronger recency effect in fast (decision or tapping) blocks as compared to slow blocks, whereas blue lines depict participants with the opposite effect.

We also hypothesized that conditions with faster movements, i.e. fast-decision and fast-tapping blocks, should be associated with a stronger prioritization of earlier information. Also contrary to that prediction, there was an even stronger *recency effect* in fast decision (*S* = 47, *p* < 10^-5^) and tapping blocks (*t* = 4.33, *p* < 10^-4^). Critically, these findings remained significant after matching experimental blocks in decision and movement -sessions in decision duration by a subsampling procedure (see Figure 3B and Materials and Methods): Also in this control analysis there was a stronger recency effect in fast-decision (*t* = 7.75, *p* < 10^-9^) and fast-tapping blocks (*S* = 39, *p* = .02). This attests that experimental conditions took influence how participants arrived at their decisions, beyond mere changes in decision duration. Furthermore, these differences in temporal weighting profiles remain significant across a wide range of ‘settings’ for deriving these profiles, highlighting the robustness of these findings (see Supplemental Material). In sum, although unexpected, results are consistent with the idea that movement characteristics alter decision making in a similar fashion as shifts in decisional speed-accuracy tradeoff do.

## Discussion

*Embodied decision-making* is a perspective which assumes an inherent functional interdependency between deciding amongst different realizable events and preparing movements corresponding to their implementation (Cisek & Pastor-Bernier, 2014; Wispinski et al., 2020; Yoo & Hayden, 2018). Our findings are in support of such interdependency with regards to the pace with which decisions and movements are completed. Faster as compared to slower decisions were followed by faster finger tapping movements, even if such behavior was not encouraged, nor adaptive in the context of the task. Likewise, faster as compared to slower tapping induced faster decision-making, which cannot be easily reconciled with strategic considerations. Furthermore, inflicting faster tapping seems to alter decision-making in a similar fashion as conditions favoring faster decisions: In both fast-tapping and fast-decision blocks, decisions relied more strongly on information presented shortly rather than long before movement onset. Taken together, these findings suggest that decisions not only determine movement characteristics, but that properties of movement constrain the time and manner with which decisions are made.

Previous studies suggest that decisions and movements are co-regulated to increase capture rate, which declines with the average time and effort required to obtain reward (Shadmehr et al., 2019). It is plausible that such coregulation is inherent to the sensorimotor system, and therefore not limited to contexts explicitly promoting such behavior. To address this hypothesis, in the present study coregulating decisions and movements was not effective to alter capture rate: Tapping speed in the movement phase did not change the pacing of a trial, nor did the duration of the decision phase, with the exception of fast-decision blocks, to encourage faster decisions. Hence, it was not adaptive in terms of capture rate to speed up decisions or finger tapping, unless this was explicitly instructed. Importantly, we did not find that understanding of these instructions modulated the experimental effect (see Supplemental Material). Moreover, tapping faster after faster decisions, as observed in the decision session, did not allow to increase the amount of momentary reward earned, as a timeout of three seconds in the movement phase ensured that the tapping sequence was virtually always completed on time (and hence rewarded). Likewise, deciding faster before fast finger tapping, as observed in the movement session, did not allow to increase the amount of momentary reward neither, as faster decisions could not increase probability of success. Finally, changes in effort allocation likely cannot account for these findings neither. As such, one may assume that faster, more effortful finger tapping (Arias et al., 2015; Jäncke et al., 2006; Lutz et al., 2004) was compensated by performing shorter, less effortful decisions (Petitet et al., 2021), and vice versa, explaining the findings in both sessions. However, by nature of the task late decisions benefitted from more conclusive information than early information, which was intentionally held ambiguous or even misleading, to prevent participants from deciding too quickly (see Materials and Methods). In line with this, drift rate was significantly reduced in fast-decision blocks as compared to slow-decision blocks, suggesting that deciding with limited evidence and under time pressure may have been more difficult, and hence more effortful, rather than less effortful. In addition, a recent study explicitly tested in a related task whether movement effort is compensated by shortening decision duration, finding no such effect with 31 participants (Lunazzi et al., 2021). Given these considerations, we interpret current findings as reflecting inherent properties of the sensorimotor system rather than strategic adaptations with changing task contexts.

If not strategically advantageous, why did the pacing of decisions and movements condition each other? Several brain areas have been proposed, which may take common influence on decision making and movement control. Subcortical structures, specifically the basal ganglia (Forstmann et al., 2008; Herz et al., 2016; Thura & Cisek, 2017) and the locus coeruleus-noradrenergic system (Hauser et al., 2018; Murphy et al., 2016; Steinemann et al., 2018), may broadly regulate activity of cortical sensorimotor areas when time pressure is high. The core idea of these proposals is that neural activity in sensorimotor cortex accumulates towards commitment for a certain motor response (Kelly et al., 2021; Murphy et al., 2016; Steinemann et al., 2018; Thura & Cisek, 2016). Under time pressure, these subcortical areas may upregulate neural activity in cortical structures so that such decisional commitment occurs earlier. On the level of cortical premotor and motor areas, such global change in neural activity can be observed as broad disinhibition of the motor system: Beta and mu power, electrophysiological markers of inhibitory control over sensorimotor sites, are reduced when participants are required to report decisions fast through movement (Kelly et al., 2021; Murphy et al., 2016; Pastötter et al., 2012; Steinemann et al., 2018). Likewise, stronger motor disinhibition at corresponding sensorimotor sites is associated with faster response times (Little et al., 2019; Meziane et al., 2015; Wessel, 2020) and higher movement velocity (Chen et al., 2007, 2011; Pogosyan et al., 2009; Torrecillos et al., 2018; but see Tatti et al., 2019). Such motor disinhibition may therefore reflect an increased tendency to execute a movement, and hence to report a decision, thereby reducing the time available to decide. Taken together, we propose that subcortical structures, such as basal ganglia or the noradrenergic system may regulate broad changes in neural activity of the sensorimotor cortex when fast action is required, reducing the time taken to decide and to move in a joint fashion. Future studies may turn to electrophysiological correlates of decisions (see Kelly & O’Connell, 2015) and movements (see Wessel, 2020), to further insight on the cortical mechanisms acting on the level of both decisional and movement control.

Such interpretation of a global signal originating from subcortical areas, controlling the urgency with which both decisions and movements are made, is in line with our findings on temporal weighting profiles. Participants prioritized recent over earlier information when making their decisions. This *recency effect* in the temporal weighting of information was enhanced in conditions favoring either fast decisions or fast tapping, suggesting that time pressure on either aspect of the task had a comparable effect on decision-making. These findings are conceptually in line with abovementioned proposals that cortical activity representing decisional evidence is dynamically amplified by subcortical projections, in order to allow to reach decisional commitment faster when the response deadline is approaching and time pressure to respond is high (Cisek et al., 2009; Thura, 2020; Thura et al., 2012). Our findings suggest that both faster decision-making, as well as faster tapping movements, may have induced such pronounced dynamic amplification, leading to an increased reliance on decisional information as the onset of movement approaches.

These findings on temporal weighting profiles provide strong evidence that not only the decision policy, but also movement characteristics such as tapping speed, constrain how people arrive at their decisions with time. However, we initially predicted a stronger *primacy effect* when fast decisions or tapping were required, i.e. the opposite of what we observed. Two aspects may account for these unexpected results. First, the task design itself may have promoted a recency effect: By nature of the task, later information was more indicative of the correct decision than earlier information, since less screen refreshes remained which could sway the trial outcome. In addition, early information was held more ambiguous to prevent participants from deciding too early (see Materials and Methods). As prioritizing late over earlier information was more advantageous in our task, participants likely adapted this behavior (Levi et al., 2018). Secondly, we predicted a stronger primacy effect in conditions promoting faster finger tapping, since increased costs of motor control should lead to a stronger gradual commitment towards a singular response. It remains possible, however, that the tendency to tap faster made it instead necessary to prepare both movements equivalently, so that either one could be executed effectively after the decision (Haith et al., 2016). From this perspective, increased motor demands may have called for less, rather than more, response selectivity. Indeed, some studies suggest that unchosen responses are prepared more when deciding and moving under time pressure, suggesting less selectivity of preparing a singular motor response (Murphy et al., 2016; Steinemann et al., 2018). In short, subcortical networks may instigate higher activity in cortical (pre-)motor areas allowing faster commitment to either response, leading to less response selectivity and ensuring that either response can be executed swiftly. This would leave greater flexibility in selecting the appropriate movement based on late decisional information, i.e. causing a stronger recency effect. However, more research is needed on this matter, as two other studies reported no change in excitability of unchosen motor representations across urgency contexts (Derosiere et al., 2021; Spieser et al., 2018).

At first glance, our findings seem to be at odds with results of Reynaud and colleagues (2020), who also investigated how movement duration affects decision-making. In contrast to our findings, faster as compared to slower movements were associated with *longer* decisions (see also Lunazzi et al., 2021). However, in their task 160 correct decisions completed an experimental block, and faster movements allowed to finish trials faster. Hence, performing slow, accurate decisions and fast movements posed the best strategy to finish early. As such, trials requiring fast movement may have allowed to invest relatively more time in deciding accurately. This was critically different from our experiment where tap duration did not change trial length and where experimental blocks had a fixed duration, independent of performance. As a result, in our study it was not advantageous to trade off decisions and movements, likely leading to the observed coherence of these sensorimotor processes. Bridging findings of both studies, properties of the sensorimotor system may promote a natural matching of the pace of decisions and movements, but contrary policies may be implemented if advantageous in the task context.

A perspective of embodied decision-making argues against a strictly sequential relationship of decisions and movements. Hence, it is somewhat a caveat that our data analysis was based on approaches assuming that reaction times reflect the additive sum of sensorimotor delays attributable to decisions and movements. Such assumption was made when subtracting simple reaction times from reaction times to isolate decision durations (see Materials and Methods), as well as when fitting reaction times to drift diffusion models. Yet, we chose to adopt such an approach to ensure that faster decisions in fast-tapping blocks could not be attributed to shorter motor delays. A mathematical formalization of behavior from an embodied decision-making perspective is still in its infancy (e.g. Lepora & Pezzulo, 2015), which will be required in the future to provide an alternative view on sensorimotor behavior.

## Conclusion

Our results demonstrate an interplay between decisional and motor processes. Movement characteristics not only depended on properties of decisions preceding them, but surprisingly, they also took distinct influence on when and how people arrived at their decisions. We interpret these behavioral results in the context of embodied decision-making, whereby shared neural mechanisms may not only enable faster movements but also assist in making decisions in less time. These findings open interesting new perspectives for future research, which may allow to reconceptualize sensorimotor processes and their associated brain regions as a functional gradient ranging from simulating potential action outcomes based on environmental information to establishing preferred action outcomes in the environment through movement.

## Materials and Methods

### Participants

The final dataset consisted of 62 right-handed participants (15 male) between 18 and 34 years old (*M* = 23.69, *SD* = 3.29), which were recruited on Facebook in groups for paid participation in academic research at Belgian universities. They affirmed to have normal or corrected-to-normal vision, no colorblindness, no history of neurological, psychiatric or mental disorders and no physical injuries or disabilities. Participants were asked to perform two sessions of the experiment circa seven days apart, of which one was the decision session and the other the movement session. Data from both sessions was available for 55 out of 62 participants, which were conducted 7.22 days apart on average (Range = 5 - 14 days, *SD* = 1.08). In the cases data was not available for one session (7 participants), the causes were due to early termination of the session, an interruption of the internet connection, a refresh rate of the computer monitor lower than 60 Hz or poor behavioral performance leading to the exclusion of more than 50 % of the trials (see Data analysis). In sum, data was available for 58 decision sessions (30 as second session) and 59 movement sessions (29 as second session). This was in line with the goal to reach 60 participants per session, in order to detect small-to-medium effect sizes (*d_z_* ≥ 0.37) in paired-samples *t*-tests, and small-to-medium correlations (*r* ≥ 0.25) with 80 % power. Apart from these 62 participants in the final dataset, eight more participants started at least one session, but were excluded due to similar reasons as stated above.

Paid 10 Euro per hour, participants received 25 Euro for circa 2.5 hours split into two experimental sessions, plus a performance-dependent bonus (see Experimental task), which was circa 5 Euro. The protocol was approved by the institutional review board of the Catholic University of Louvain, Brussels, Belgium, and was in compliance with the principles of the Declaration of Helsinki. Written, informed consent was given by emailing photographed and hand-signed consent forms to the experimenter.

### Experimental task

As seen in Figure 1, a trial started with a light gray fixation dot on a dark gray background, shown for 100-500 ms. In the subsequent pre-decision phase, two ‘baby bananas’ appeared on the left and right of the fixation dot at a 30-degree angle and covered a circular track of 60 degrees, crossing each other, to arrive centrally below the fixation dot, separated by a horizontal gap. With a fixed duration of 1000 ms, this pre-decision phase telegraphed the onset of the decision phase. In this next phase, bananas grew progressively, with their tips following a circular track around the fixation dot, whereas their ends remained in place. With every refresh of the screen, i.e. at a rate of 60 Hz, pseudo-randomly either the left or right banana was extended by 2 degrees, creating the illusion of continuous growth at a non-constant speed. If uninterrupted by the participant’s response, the bananas kept growing for 179 frames or 2983 ms (hereafter: 3000 ms) to cover 358 degrees around the fixation dot. An uneven number of frames ensured that one banana was always longer by the end of a trial. Participants had to select the anticipated longer banana before the end of the decision phase by pressing the S-key (left banana) or L-key (right banana) with their respective index finger. With this first finger tap, the movement phase was initiated by placing a rose caterpillar below the central fixation dot on a circular track parallel to the one of the chosen banana, which itself faded into the background within 200 ms. The caterpillar extended with each key press, including the initial decision, by 36 degrees along the circular track, with movement animations lasting circa 80 ms each (5 frames). In addition to the initial decision, participants thus had to tap four more times to move the caterpillar head to a position above the fixation dot. Important for the purpose of the study, the movement phase had a fixed duration of 3000 ms, i.e. the subsequent phase did not begin earlier even if participants completed the movements earlier (cf. Morel et al., 2017). To represent this fixed duration of the movement phase visually, the fixation dot disappeared with phase onset and gradually returned to initial opacity by phase offset. In addition to the caterpillar which remained onscreen, bananas additionally reappeared within the last 200 ms of the movement phase. If participants had not responded within 3000 ms during the decision phase earlier, the caterpillar did not appear, however participants still had to sit out the movement phase. Next, as a post-decision phase, bananas completed their growth paths in a similar fashion as the decision phase until they covered an approximately full circle around the fixation dot. In this way, for all experimental conditions except one (see the following paragraph), bananas grew for a fixed duration of 3000 ms on every trial independent of the timing of participant’s decision. In a feedback phase lasting 800 ms, the outline of the chosen banana was highlighted in either green or red, depending on the correctness of their decision. If participants had not responded within the specified time of either the decision or movement phase, the fixation dot was replaced with the caption ‘decide faster!’ or ‘tap faster!’, respectively. A final intertrial phase of 700 to 1000 ms followed, which was visually identical to the fixation phase.

Trials were embedded in different blocks, which manipulated either decision speed or tapping speed experimentally, depending on the session: In the decision session, blocks encouraged either fast decisions or slow (accurate) decisions, whereas in the movement session, either fast or slow finger tapping was required. Blocks in the decision session differed in that fast-decision blocks exceptionally allowed to shorten the duration of a trial by deciding faster: In contrast to all other conditions (see Figure 1), bananas grew four times faster in the post-decision phase (by only showing every fourth frame of growing bananas). Additionally, fast-decision blocks were not limited to 40 trials as in other blocks, but instead the length of these blocks was fixed to 5 minutes and 30 seconds. Participants were instructed to select as many bananas as possible and were informed that choosing bananas quickly would allow them to complete more trials. To encourage this behavior, a block-based bonus of 10 cents was given when participants ‘completed more rounds than average’, i.e. completing more than 40 trials in fast-decision blocks. In contrast, for slow-decision blocks, participants were informed that exactly 40 trials had to be completed, with a fixed trial duration independent of behavior. Here, participants had to select the correct banana as often as possible. This was again encouraged by rewarding an ‘above average’ amount of correctly chosen bananas with a block-based bonus of 10 cents, i.e. when more than at least 85% of the decisions were correct. Importantly, it was stressed that tapping speed in the movement phase would not alter trial duration in any experimental condition. This fact was further confirmed with the visual representation of an example trial, which showed that deciding faster in fast- decision blocks was the only circumstance in which behavior would change trial duration.

Blocks in the movement session differed in that fast-tapping blocks required to move the caterpillar with a minimum speed, whereas slow-tapping blocks were limited by maximum speed. For this purpose, a snake either chased the caterpillar during the movement phase or retreated from it, both at a constant speed calibrated individually per participant (see Session Protocol). In fast-tapping blocks, the snake was initially placed 36 degrees behind the caterpillar and moved at a uniform velocity corresponding to 85 % of maximum tapping speed. In slow-tapping blocks, the snake was initially placed 36 degrees in front of the caterpillar and retreated with a constant pace corresponding to 59.5 % of maximum tapping speed. Depending on the condition, the snake’s color was either purple or green, counterbalanced between participants. If participants tapped too slowly or too quickly, respectively, the caterpillars head would coincide with the snake’s head and both animals would stop moving. In this case, novel responses would no longer be registered, otherwise the trial continued as described earlier. Also for the movement session an example trial demonstrated that trial duration was fixed and thus independent of the participants’ behavior.

In each session, participants performed four times each of the two block types, i.e. eight blocks of (at least) 40 trials presented in randomized order, resulting in (at least) 160 trials per experimental condition. Each block started with a fixed instruction screen of at least 20 seconds before moving on, ensuring that instructions for the following block could not be skipped. Each first trial of a block had an additional 700 ms of fixation. To further remind participants of the current instructions in each block, a symbol representing each of the four block types was shown in the upper right corner of the screen throughout (see Figure 1). To keep participants engaged throughout the task, 1 point could be earned for each correctly performed decision and movement, respectively, i.e. a maximum of 2 points per trial. After each block, the total number of accumulated points was converted to a monetary bonus at a rate of 0.4 cent per point, in addition to feedback on the possible block-based bonus of 10 cent in decision sessions. Between blocks, participants were further informed about the number of remaining blocks of the current session.

Trial difficulty was controlled between experimental conditions by adjusting the rate at which the winning banana outgrew the other over time: A more asymmetric growth results in an easier decision. We intermixed four trial types which were chosen in analogy to similar approaches in the literature (e.g. Derosiere et al., 2019; Reynaud et al., 2020; Thura, 2020 in humans or Thura & Cisek, 2014, 2016, 2017 in monkeys): In ‘obvious’ trials, one banana grew larger than the other one early on and remained so for the remainder of the trial. In ‘ambiguous’ trials, bananas remained in competition for length for about halfway of the trial, before one banana proceeded to take the lead. In ‘misleading’ trials, one banana seemed to outgrow the other early on in the trial but lost that competition by the end of the trial. Finally, in ‘random’ trials bananas could show any growing pattern. Each experimental condition consisted of 30 % obvious, 30 % ambiguous, 20 % misleading and 20 % random trials.

### Session outline

The online experiment was built in lab.js, hosted on Open-Lab (Henninger et al., 2019), and was accessed via the Google Chrome browser. It required a screen of at least 15 inch in diameter with a refresh rate of 60 Hz, and was run in full screen mode. At the start of each session, participants were asked to create a calm environment by using a solitary room and by removing possible sources of noise. An overview of the session outline can be seen in Supplemental Figure S5.

The movement session always started with a so-called ‘keyboard calibration’, which was skipped for the decision session. This segment allowed to estimate maximum tapping speed, although this goal was not disclosed. Within four-second intervals, the S- or L-key had to be tapped repeatedly as fast as possible. Each trial consisted of a countdown represented as a pie chart, which declined from 360 degrees (full circle) to 0 degrees (no circle) within four seconds. The letter which had to be tapped was superimposed. A counter for the total number of key taps was shown below the pie chart, both were reset with each new trial. After each trial, the feedback to tap ‘FASTER!’ was given irrespective of performance. Participants performed eight trials per key (the beginning key was randomized) and the maximum tapping speed was calculated as average tapping speed for the last six trials of both keys.

The first out of two sessions continued with a familiarization segment. This segment introduced the decisional aspect of the task through visual animations and written text. Instructions could be repeated at will by navigating through instruction screens with dedicated keys, however new instructions could not be skipped. A simpler version of the task with only the decision phase could then be practiced in 20 trials, i.e. a trial required only one key press to report the decision. Practice had to be completed up to three times if the proportion of correct decisions did not reach at least 70%. In that case, participants were given the choice to re-read or skip instructions before repeating practice trials. All sessions then continued with a training phase, which introduced the movement phase of the trial. Participants were then informed about the two experimental blocks in the session. This full task had to be practiced in another 20 trials, with 10 trials in each condition. Training had to be again completed up three times if either a decision accuracy of at least 70% was not reached, or the required number of key taps was below 70%.

The main experiment followed with circa 320 trials. Participant’s understanding of the instructions was afterwards probed with a questionnaire: Statements on the influence of behavior on trial duration had to rated on agreement (see Supplemental Material). Participants with a good task understanding should indicate that choosing a banana faster or moving the caterpillar faster does not alter trial duration, except for fast decisions in fast-decision blocks.

Finally, participants performed a simple reaction time (SRT) task, which was comparable to the main task, but stripped of decisional aspects by giving away the correct response in advance. This task allowed to estimate sensorimotor delays attributable to pressing the first key of a tapping sequence (see Data analysis and Supplemental Material). In the pre-decision phase of the SRT task, one motionless baby banana was pointing towards the left or right, randomized between trials. After a delay of 1000 to 1600 ms, a fully extended banana covered 180 degrees in the same direction. As soon as the banana thus fully grew, the corresponding key had to be tapped. Premature responses were followed with the words ‘too early!’ for three seconds instead of the extended banana. All other phases of the trial were identical to the main task, with the exception that there was no post-decision phase. The movement session entailed 40 SRT trials requiring fast tapping (promoted by a chasing snake) and 40 SRT trials requiring slow tapping (promoted by a retreating snake; with block order randomized). The decision session entailed 40 SRT trials with no restrictions on tapping speed (with no snake), as was the case for the main task.

At the end of the second session, participants had to fill in the S-UPPS-P questionnaire (Cyders et al., 2014). This impulsivity questionnaire was administered to potentially account for individual differences in the time allocated for choosing and acting. Since such individual differences in impulsivity were not central to the purpose of this study, these data are not reported.

### Data analysis

#### Behavior

Endpoint measures were computed which reflect decision and motor performance: *Decision duration* was computed for each participant as average reaction times in each experimental condition minus the average reaction time in the corresponding SRT block (c.f. Derosiere et al., 2019; Reynaud et al., 2020; Lunazzi et al., 2021; Thura, 2020; Thura & Cisek, 2014, 2016, 2017). This ensured that only the time taken to decide was compared between experimental blocks, without confounding it with the time needed to perform a first finger tap. *Success probability* reflects the objective probability that the decision is correct, given the length difference in bananas at the time of the decision (see Supplemental Material for further details on this measure). *Decision accuracy*, as a related measure, was calculated as the factual proportion of trials in which the chosen banana won. *Tap duration* was measured as average time difference between successive finger taps and hence reflects how fast participants moved the caterpillar in the movement phase of the task. *Movement accuracy* reflects the proportion of trials in which the four finger taps were carried out at the proper pace in the movement phase.

Trials with no response during the decision phase were removed from both the main experiment and SRT task. Trials with anticipation errors were also excluded: These regarded key presses occurring in the pre-decision phase, decisions with a duration less than 150 ms in case of the main task, or with a reaction time less than 50 ms in case of the SRT task. Trials of the main experiment were excluded if frame rate deviated by at least 1 Hz from the desired 60 Hz in either pre-decision, decision or movement phases. For the SRT task frame rate was not recorded. *Decision duration* was calculated based on trials with correct decisions, i.e. the chosen banana was the correct one, and with correct movements, i.e. the four finger taps were carried out at the proper pace within the movement phase. *Success probability* and *decision accuracy* were calculated based on trials with correct movements. *Movement accuracy* was calculated based on trials with correct decisions. Overall, 22% (decision duration), 8% (success probability and decision accuracy) or 20% (movement accuracy) of trials were thus removed from the main experiment, and 7% of trials were removed from the SRT task.

Statistical tests were conducted separately for each session, which allowed more targeted testing of the hypotheses. Robust statistical methods were used to address apparent issues with non-normality and heteroscedasticity (see Figure 2): To compare two measurements of the same participants, signed-rank tests were used. For more than two measurements, (Greenhouse-Geisser corrected) repeated-measures ANOVAs were used. To test ANOVAs for robustness, results were compared for consistency before and after removing participants with extreme values. Extreme values were defined as data points lying outside of three median absolute deviations of the median of all cells of the corresponding ANOVA (Leys et al., 2013). Pairwise comparisons following ANOVAs were again conducted with signed-rank tests and corrected for multiple comparisons by the method of Bonferroni-Holm (Chen et al., 2017). Linear relationships were tested using weighted-least squares regression. In this approach, the weight of individual data points on the regression slope is inversely proportional to the increase in spread of these data points with changes in the predictor variable, thereby accounting for heteroscedacity. These regressions were further tested for robustness by repeating the same test after removing outliers. Outliers were defined as data points with a Cook’s Distance three times larger than the group mean (Cook, 1977). Statistical analysis was conducted in R (R Core Team, 2018), relying on packages BDSA, rstatix and stats.

#### Drift diffusion modeling

To corroborate the interpretation of behavioral data as reflecting shifts in duration with which decisions and finger tap movements are performed, drift diffusion models were fit with the hddm- package in Python (version 0.7.3; Wiecki, 2013). Four parameters characterize different aspects of the sensorimotor process: The decision threshold *a* reflects the decisional speed-accuracy tradeoff with higher values indicating that more certainty is needed before commitment, resulting in slower but more accurate decisions. The drift rate *v* reflects the efficiency with which a correct decision can be made. It therefore increases with lower task difficulty, and higher motivation and performance. The non-decision time *t* represents sensorimotor delays unrelated to decision-making, such as sensation and response execution. The response bias *z* reflects an a-priori tendency to prefer one response over the other, for example when one response is more rewarded than the other.

In line with our hypothesis, for each session a model allowing the decision threshold and non-decision time to change with experimental conditions was fit. This model was compared with models of different complexity (see Table S2). The most complex model assumed that, in addition to the decision threshold and non-decision time, also the drift rate differed between experimental conditions. All other models assumed a fixed drift rate, as well as at least one of the parameters of interest (decision threshold and non-decision time) to be fixed between experimental conditions. For all fitted models, we assumed no response bias since an equal amount of left and rightward responses was required in the task (i.e. *z* was assumed to be 0.5, see e.g. Assink et al., 2015; Herz et al., 2016). Although this may be a simplifying assumption, it allows for a more powerful test of the effects of interest (Lerche & Voss, 2016; van Ravenzwaaij et al., 2017). Moreover, our behavioral findings likely cannot be accounted by response biases. For each model, ‘flexible’ parameters were modelled as within-subject effects by fitting one parameter value to the slow condition of the respective session, and another parameter value reflected the deviation thereof in the fast condition (see Table S2). Models were hierarchical in that parameter values of each subject were assumed to spread around the group mean of that parameter. Correct and incorrect responses were taken as upper and lower boundaries of the drift diffusion process. Five percent of responses were assumed to be outliers, as recommended to improve model fit (Wiecki et al., 2013).

Models were fit to reaction times of the decision phase of the task (see Figure 1), i.e. the time from stimulus onset to the first key press in a trial. This allowed to test whether faster finger tapping in the movement phase of the task was associated with a faster initial finger tap in the decision phase. Modelling further provided an alternative way of estimating (non-)decision time, without the SRT task, to ensure that presented findings are consistent across different methodological approaches. Consistent with this aim, reaction times less than 150 ms were discarded as anticipation errors, rather than decision durations below 150 ms as was the case in the behavioral analysis. Each model in Table S2 was fit twice with 5000 discarded samples as burn-in and keeping every third of 6000 subsequent samples, resulting in 4000 final posterior samples per parameter. Model convergence was assessed by visual inspection of the Markov-chain traces, as well as computation of the Gelman-Rubin statistic. With values below 1.1 considered satisfactory by convention (Brooks & Gelman, 1998), Gelman-Rubin values indicated convergence for all models of the decision session (all below 1.04), as well as the movement session (all below 1.01). Models were then compared based on the bias-corrected Bayesian Predictive Information Criterion (BPIC, Ando, 2007; Heathcote et al., 2015), since this criterion does not require the true model to be amongst candidate models. Best-fitting models were further ratified by posterior predictive checks: New data was simulated from best-fitting models and visually compared to the real empirical data (see Figure S6), suggesting good agreement between model predictions and reality. As a final step in this model analysis, for the best-fitting models, the probability of the existence of within-subject effects was expressed as proportion of posterior samples smaller than zero for the respective parameter. This proportion reflects the estimated probability that parameter values are more negative in fast conditions than in slow conditions. In contrast to *p*-values, a probability exceeding 97% indicates that an effect likely exists (Makowski et al., 2019).

#### Temporal weighting profiles

To test the predictions that movement preparation induces a gradual commitment towards a decision, and more so when fast movement is required, temporal weighting profiles were computed. These profiles outline in how far newly presented momentary evidence is predictive of the subsequent decision, depending on when this evidence is presented relative to movement onset. As a first step in computing these profiles, momentary evidence presented throughout a trial was quantified: In time windows of 200 ms, frames shifting evidence in favor of either the left (−1) or right (+1) banana were summed up to a total value. As such, momentary evidence could either be positive or negative, with larger values indicating stronger momentary evidence towards the left or right. Momentary evidence was summarized in this way in a moving 200-ms window across the entire sequence of decisional evidence presented throughout a trial. This approach is in line with the notion that sensory evidence is likely integrated into decisions at a rate of circa 5-6 Hz in humans (Carland et al., 2015; Cisek et al., 2009; Scott, 2016). Momentary evidence and decisions (−1 for left and +1 for right responses) were then correlated across trials for each participant, experimental condition and time window of presentation (see Bitzer et al., 2020, for a similar approach). As seen in Figure 3, resulting point-biserial correlations characterize in how far momentary evidence predicts decisions, depending on the time of presentation relative to estimated movement onset. To assess the relative weight of earlier versus later evidence on decisions, for each participant and experimental condition, the slopes of temporal weighting profiles were extracted for the estimated decision duration, i.e. negative median decision duration to zero (see Figure 3, see Levi et al., 2018; Spieser et al., 2018, for similar approaches). Negative slopes suggest that earlier evidence is more predictive of decisions than later evidence (*primacy effect*), whereas positive slopes suggest the opposite (*recency effect*). For each experimental session, slopes were then compared with signed-rank tests, or with paired *t*-tests in case Shapiro–Wilk tests were not violated (see Hubert-Wallander & Boynton, 2015; Levi et al., 2018, for similar approaches).

To ensure that main results were not driven by differences in average decision duration between experimental conditions (Okazawa et al., 2018), for a control analysis we applied a subsampling procedure: Decision durations were first grouped into time bins of 100 ms. Per participant, trial numbers in each time bin were then matched between compared experimental conditions by subsampling (see Murphy et al., 2016, for a similar approach). The remaining trials, now matched in decision duration, were then again used to calculate temporal weighting profiles. Resulting temporal weighting profiles were averaged across ten iterations of such subsampling before extracting individual slopes. Three participants of the decision session were removed from this control analysis since less than ten trials remained after matching decision durations by subsampling.

## Acknowledgements

Thomas Carsten is a Postdoctoral Researcher of the Fonds de la Recherche Scientifique – FNRS. This work was also supported by the Fonds de la Recherche Scientifique – FNRS under T.0082.19, awarded to Julie Duque. We want to thank Fanny Montulet and Merlin Somville for their useful comments on the methodology of this research project.

## SUPPLEMENTAL MATERIAL

### Control analyses on temporal weighting profiles

To check for robustness of reported results on temporal weighting profiles in the manuscript, additional tests were conducted. First, we repeated the main analysis after summarizing momentary evidence within time windows of 300 ms and 100 ms. As similar results were obtained (see Figure S1, 1^st^ and 2^nd^ row, all *p*’s < .01), we concluded that the chosen time window of 200 ms did not affect results critically. Furthermore, we tested whether results subsist when only trials with fully random evidence are used to compute temporal weighting profiles. Indeed, if evidence is not fully random (such as in obvious, ambiguous or misleading trials used in the experiment), earlier evidence may be predictive of later evidence and vice versa, which may prohibit to isolate the relative contribution of individual pieces of evidence on decisions (Okazawa et al., 2018). As seen in Figure S1, third row, an analysis with only random evidence -trials led to similar results (both *p*’s < .05); it also showed overall comparable shapes of temporal weighting profiles, suggesting that main findings did not critically depend on the type of trials entered in the analysis. We then tested whether findings remain significant after matching experimental conditions in each session in success probability: This test ensured that changes in temporal weighting profiles do not primarily reflect changes in overall task performance between conditions. For this purpose, a similar matching procedure was utilized as described in the Materials and Methods of the main text, only that trials were matched by success probability, not decision duration. Also in this analysis findings were significant for both sessions (see Figure S1, fourth row, both *p*’s < .01).

Next, we sought further support for the idea that an increase in tapping speed may have led to a stronger recency effect in temporal weighting profiles of decisions. Therefore, we proceeded to sort all trials as fast-tapping and slow-tapping trials, depending on whether tapping speed in that trial was exceeding median tapping speed of the respective participant and session (i.e. performing a ‘median-split’ based on tapping speed). These newly assigned fast-tapping and slow-tapping trials were then again matched in decision durations by the subsampling procedure described in the main text. As a result, for each experimental session, we compared trials with relative fast and slow tapping, which were however matched in decision durations. Again, we found that relatively fast tapping was related to a stronger recency effect than relatively slow tapping for both sessions (see Figure S1, fifth row, all *p*’s < .05), supporting our conclusions. Note that for the movement session this analysis was very similar to results presented in Figure 3 since relatively faster and slower tapping was prevalent in fast and slow tapping-blocks, respectively.

As another check for consistency, temporal weighting profiles were derived in relation to the onset of the decision phase (i.e. ‘stimulus-locked’). For all previous analyses, profiles were computed relative to estimated onset of the first finger tap (i.e. ‘response- locked’), in line with our research hypothesis. However, findings should be consistent across both types of analyses. A stimulus-locked analysis was again conducted based on conditions with matched decision durations to ensure that latencies between stimulus-onset and decisions were matched. As can be seen in Figure S1, sixth row, results were consistent with response-locked analyses (all *p*’s < 10^-5^), although the shape of temporal weighting profiles was different due to the nature of this stimulus-locked approach. Finally, Figure S1, final row, shows a control analysis in which decisional information was not summarized as momentary evidence in a moving time window of 200 ms (see main text), but as the absolute evidence shown up until the time point in question. In this analysis, evidence at e.g. -400 ms reflected the absolute length difference of bananas at -400 ms, rather than the *change in length difference* that occurred in the time window from -300 to -500 ms (i.e. momentary evidence). We again matched decision durations between experimental conditions in this analysis. Similar results were obtained (both *p*’s < .01).

In summary, additional analyses on temporal weighting profiles consistently support conclusions drawn in the main manuscript, demonstrating the robustness of these findings. They demonstrate that the idea, that shifts in recency effects were driven by changes in tapping speed, is plausible. This strengthens the idea that movement characteristics take influence on when and how participants arrive at their decisions.

### Behavior in the Simple Reaction Time task

Participants performed a simple reaction time (SRT) task, which was comparable to the main task, but stripped of decisional aspects. This allowed to estimate the time needed to perform a first key tap in order to isolate decision duration in the main task. The decision session entailed 40 SRT trials with no restrictions on tapping speed (‘free’ in Figure S2, no snake in the movement phase), whereas the movement session entailed two times 40 SRT trials requiring fast (chasing snake) or slow tapping (retreating snake). The effect of tapping requirements on tap duration and reaction times was tested in two separate repeated-measures ANOVAs for trimmed means (as more robust alternative to a conventional one-way repeated-measures ANOVA).

As intended, tap duration in the SRT task changed with tapping requirements (*F*(1.61, 51.44) = 394.83 and *p* < 10^-17^). As seen in Figure S2A, tap duration was shortest when fast tapping was required (*median* = 139 ms, *MAD* = 20 ms), followed by free tapping (*median* = 160 ms, *MAD* = 31 ms) and longest for slow tapping (*median* = 255 ms, *MAD* = 40 ms, all *p*’s < 10^-9^). As seen in Figure S2B, reaction time also varied as a function of tapping requirements (*F*(1.86, 59.59) = 30.32 and *p* < 10^-8^). Reaction time was shorter when fast tapping (*median* = 365 ms, *MAD* = 92 ms) was required as compared to slow tapping (*median* = 433 ms, *MAD* = 116 ms, *p* = .001). Interestingly, reaction time was shortest for free tapping (*median* = 315 ms, *MAD* = 71 ms. both *p*’s < 10^-6^).

Taken together, participants followed movement instructions well in the SRT task. Results further suggest that finger tapping with restrictions on the pacing (i.e. ‘fast’ or ‘slow’) takes more time to prepare than finger tapping with no speed requirements (i.e. ‘free’).

### Testing whether instruction comprehension affected behavior

Participants were instructed and demonstrated that the task was designed in a way that the speed with which decisions and finger tapping movements were completed was inconsequential for the duration of a trial, except for decision duration in fast-decision blocks, which shortened trial duration. This task design ensured that coregulating the pacing of decisions and movements was not profitable in terms of capture rate (see Discussion in the main text). To probe whether participants understood this task design, after each session a short instruction questionnaire was administered after the experimental task (see Table S1). In the decision session, one statement read: “In ‘fast-decision segments’, I felt like I could end a round earlier by moving the caterpillar faster.” This statement was critical since faster finger tapping was observed in fast-decision blocks, although only fast decisions could shorten trial duration, allowing to perform more trials. Therefore, a proper understanding of the task design should lead participants to disagree with this statement. Surprisingly, as majority, 37 participants agreed with the statement, suggesting that deciding faster was associated with a subjective feeling of saving time through movement. Fifteen participants correctly disagreed with this statement, whereas six participants did not know. More importantly however, all participants, i.e. those who (somewhat or strongly) agreed (*median* = 18 ms, *S* = 1, *p* < 10^-8^), (somewhat or strongly) disagreed (*median* = 14 ms, *S* = 0, *p* < .001) or did not know (*median* = 10 ms, *S* = 0, *p* = .03), tapped faster in fast-decision blocks, with no significant difference between the three groups (*F*(2, 10.98) = 3.45, *p* = .07), as suggested by an ANOVA based on trimmed means with INSTRUCTION AGREEMENT as between-subjects factor and BLOCK TYPE as within-subjects factor. This suggests that deciding faster may have led to a subjective feeling of saving time by tapping faster in most participants. However, instruction comprehension was not a critical factor for the experimental effect observed in the decision session.

For the movement session, proper instruction comprehension was less of a concern as movement requirements were clearly constrained to the movement phase by visualizing them as retreating or chasing snakes (see Figure 1). Nevertheless, also here we probed understanding of the task design to test whether it modulated the experimental effect. The statement “In segments in which the snake was chasing me, I felt like I could end a round earlier by choosing the banana faster.” was critical since participants performed faster decisions in these types of trials although this could not increase capture rate (see Discussion in the main text). A proper understanding of the task design should therefore lead participants to disagree with this statement. Twentynine participants correctly disagreed, 21 falsely agreed and nine did not know. More importantly however, faster decisions in fast-tapping blocks were observed in those participants who correctly disagreed (*median* = 84 ms, *S* = 4, *p* < .001), whereas there was no significant effect for those participants who disagreed (*median* = 21 ms, *S* = 7, *p* = .36) or did not know (*median* = 66 ms, *S* = 2, *p* = .36). However, there was no significant INSTRUCTION AGREEMENT x BLOCK TYPE interaction (*F*(2, 19.82) = 0.99, *p* = .39), indicating that there was no significant difference in effect size between groups. This suggests that fast finger tapping may have led to a subjective feeling of saving time by deciding faster in a minority of participants. However, instruction comprehension was not a critical factor for the experimental effect observed in the movement session, as a clear experimental effect was observed for the majority of participants who had a good comprehension of the task design.

### How success probability of decisions is computed

As an objective measure of the quality of decisions, success probability was computed. In each trial of our task, evidence continued to change dynamically after the decision. Therefore, decisions had a probability of being correct, corresponding to the objective probability that the chosen banana would be longest by the end of the trial. Therefore, success probability of each decision can be formally calculated as the proportion of all hypothetical growth patterns of bananas following the decision in which the chosen banana wins. In accordance with previous work (Derosiere et al., 2019; Reynaud et al., 2020; Saleri Lunazzi et al., 2021; Thura & Cisek, 2014, 2016, 2017), success probability can therefore be computed as

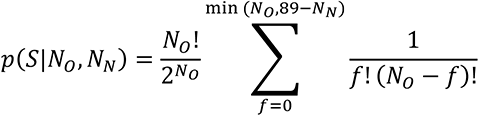

where the probability *p* of the selected banana *S* winning is equivalent to the probability of unselected banana *N* not winning. *N_N_* is the number of frames in favor of the unselected banana, *N_O_* is the number of outstanding frames, both at the estimated time of decision offset (i.e. ‘decision duration’). The selected banana *S* wins if the number of frames in favor of the unselected banana *N* stay below 89, i.e. less than half of the 179 frames. The success probability of the selected banana therefore depends on the total number of possible growth patterns for the remaining frames *N_O_*, in which the number of frames in favor of the unselected banana remain below 89, relative to the total number of all hypothetically possible growth patterns for these frames.

**Figure S1.**
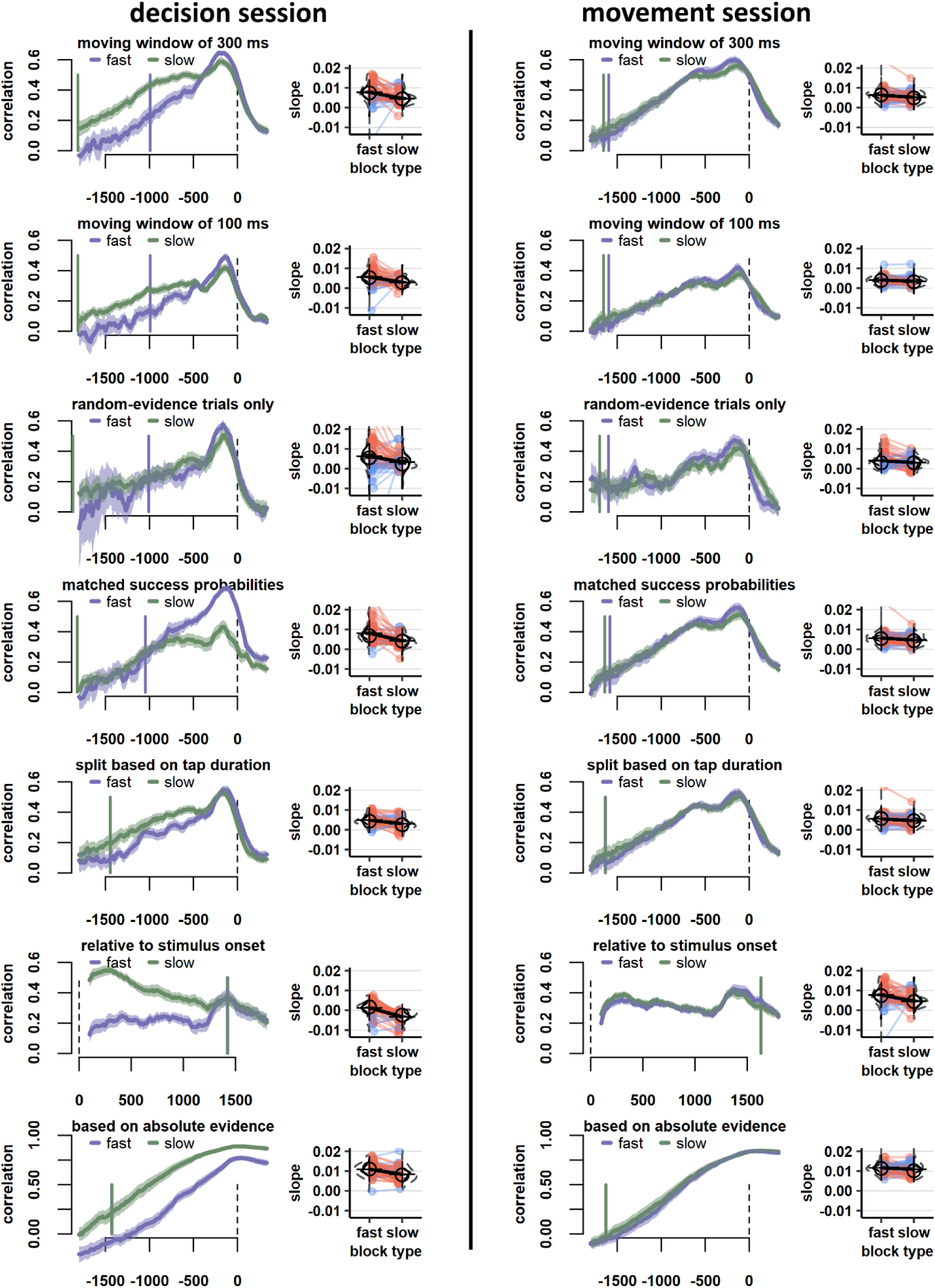
Additional analysis of temporal weighting profiles. Results were consistent across all additional analyses on temporal weighting profiles. There was a stronger recency effect in temporal weighting profiles in fast (decision or tapping) blocks as compared to slow blocks when summarizing momentary evidence in time windows of 300 ms (1^st^ row), summarizing momentary evidence in time windows of 100 ms (2^nd^ row), including only trials with fully random evidence (3^rd^ row), matching success probabilities between experimental conditions with a subsampling procedure (4^th^ row), splitting trials into relatively fast and slow tapping- trials and again matching these new ‘conditions’ for decision durations (5^th^ row), computing temporal weighting profiles relative to the onset of the decision phase and again matching conditions for decision durations (i.e. stimulus-locked, 6^th^ row), and computing temporal weighting profiles based on absolute rather than momentary evidence, again matched for decision durations (final row). For more elaboration on visual aspects of these figures, consult the caption of Figure 3 in the main manuscript.

**Figure S2.**
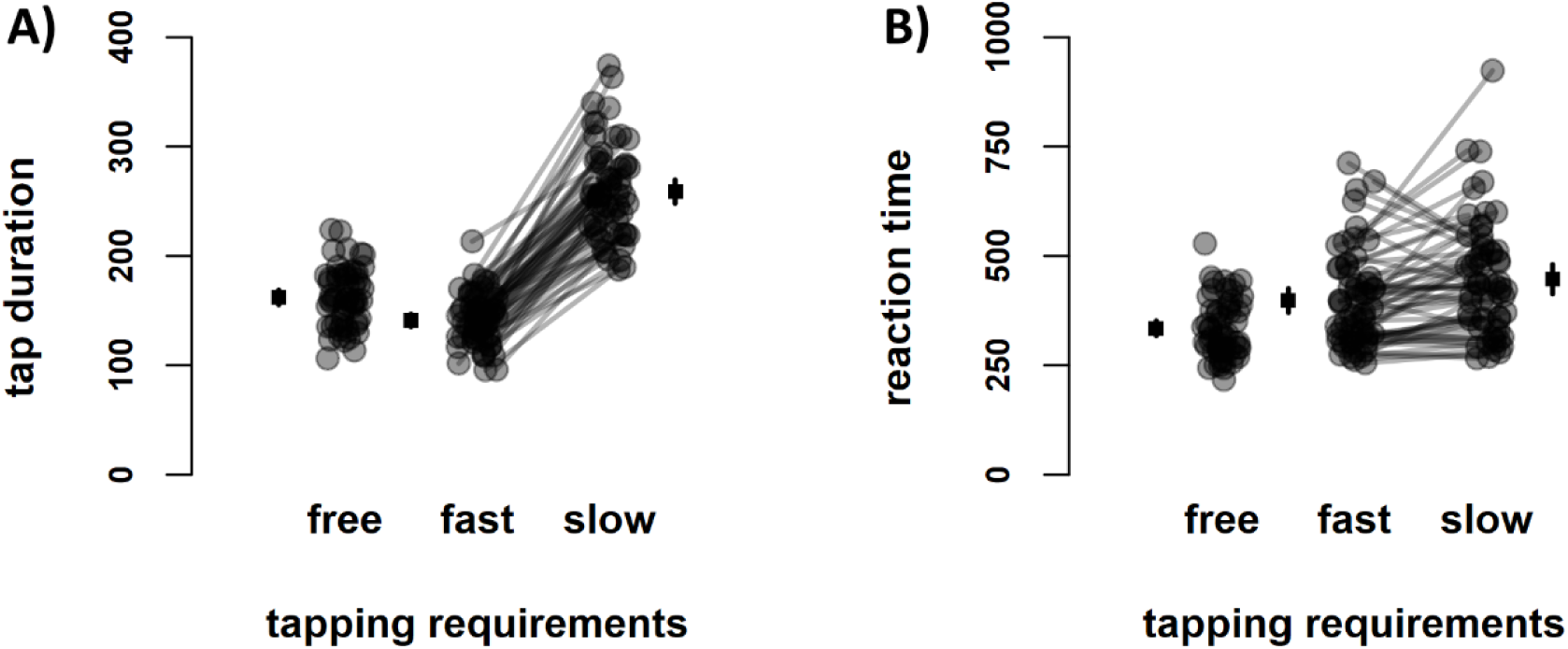
Behavior in the simple reaction time task. A simple reaction time task with no decisional aspects was administered, in which participants were required to tap with no speed constraints (‘free’, in decision session), or fast or slowly (in movement session). **A)** Tap duration was shortest when fast tapping was required, followed by free-tapping and slow- tapping blocks. **B)** Simple reaction times were fastest when no requirements on tapping speed were given, followed by fast-tapping and slow-tapping blocks. **A), B)** Individual data points are shown, with data stemming from the movement session linked with a line. Individual data points (circles) are flanked by group means (squares). Error bars of group means reflect 95%- Confidence Intervals.

**Figure S3.**
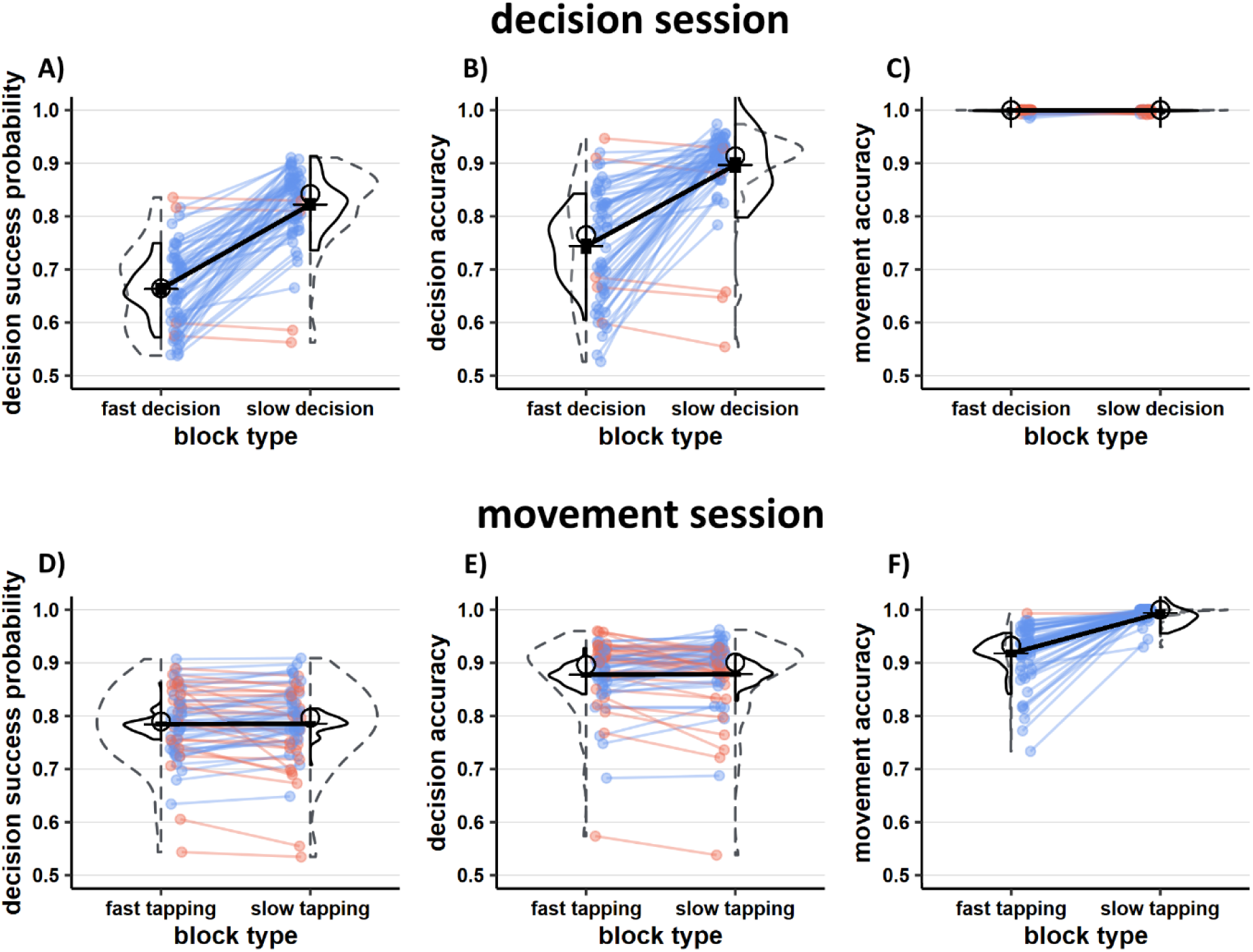
Supplemental behavioral results. **A)** Probability that decisions led to successful outcomes was significantly decreased in fast-decision as compared to slow-decision blocks. **B)** As consequence, the proportion of correct decisions was significantly reduced in fast-decision blocks. **C)** In the decision session, participants absolved virtually all tapping movements correctly, with no significant difference between blocks. **D)** There was no significant change in decisional success probability from slow-tapping to fast-tapping blocks. **E)** Likewise, there was no significant change in the proportion of correct decisions between blocks of the movement session. **F)** Participants performed significant less fast-tapping movements correctly than slow-tapping movements.

**Figure S4.**
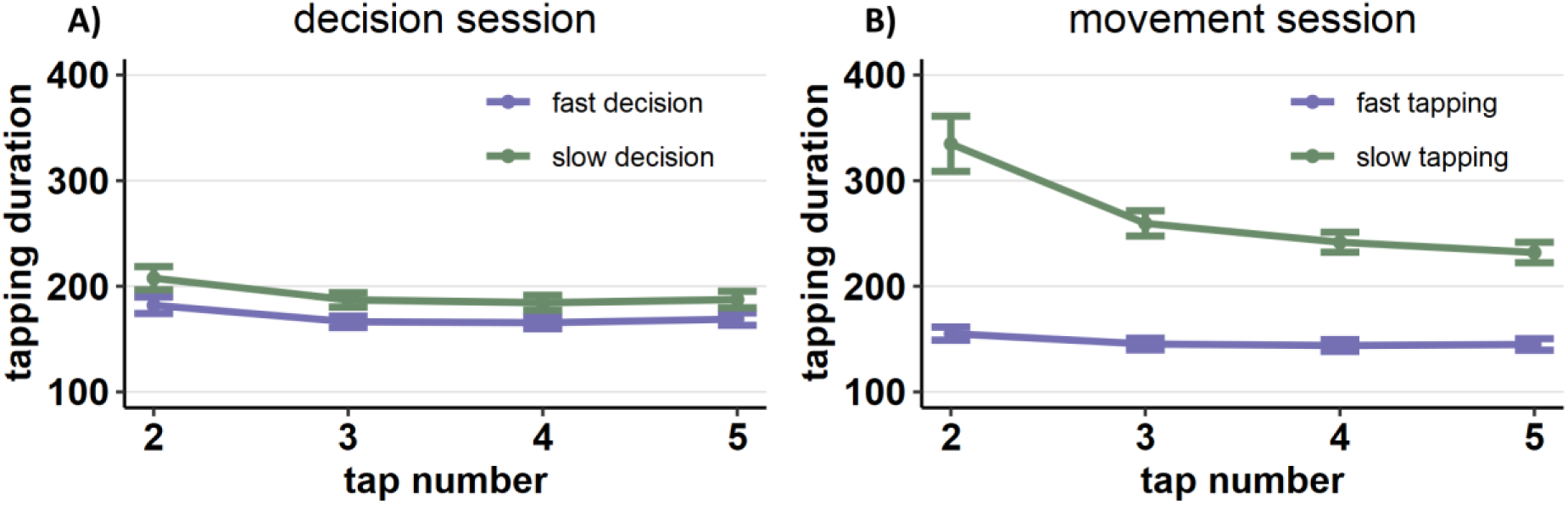
Average duration per finger tap in the movement phase for each experimental condition. **A)** In the decision session, each tap in the movement phase was faster when fast decisions as compared to slow decisions were required, demonstrating the anticipated experimental effect. **B)** In the movement session, each tap in the movement phase was faster when fast tapping as compared to slow decisions were required, demonstrating that the experimental manipulation was successful. Note that the first finger tap in each trial reflected the reporting of the decision in the decision phase, hence for this analysis 2^nd^ to 5^th^ finger tap are considered. Error bars reflect 95%- confidence intervals.

**Figure S5.**
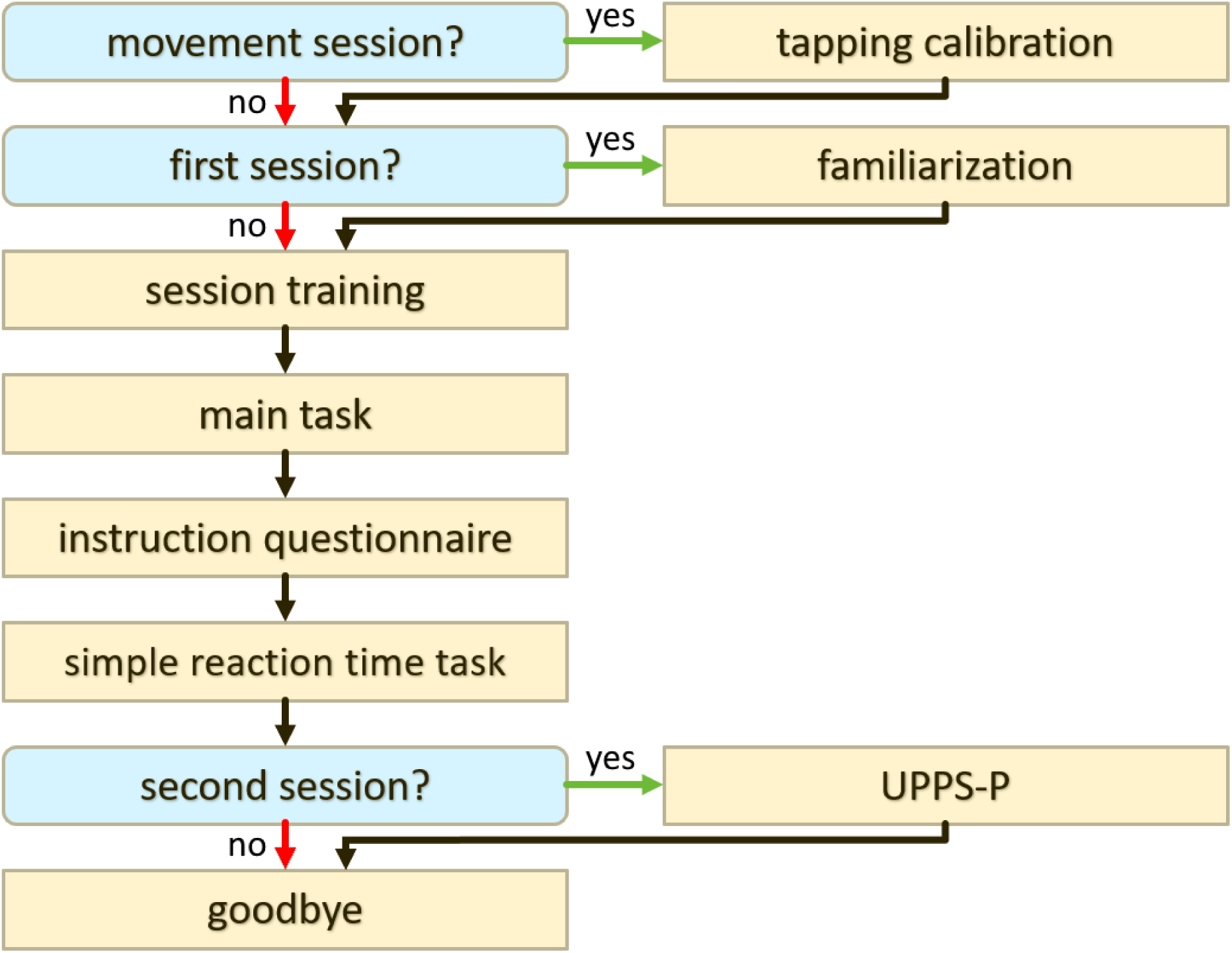
Session outline.

**Figure S6.**
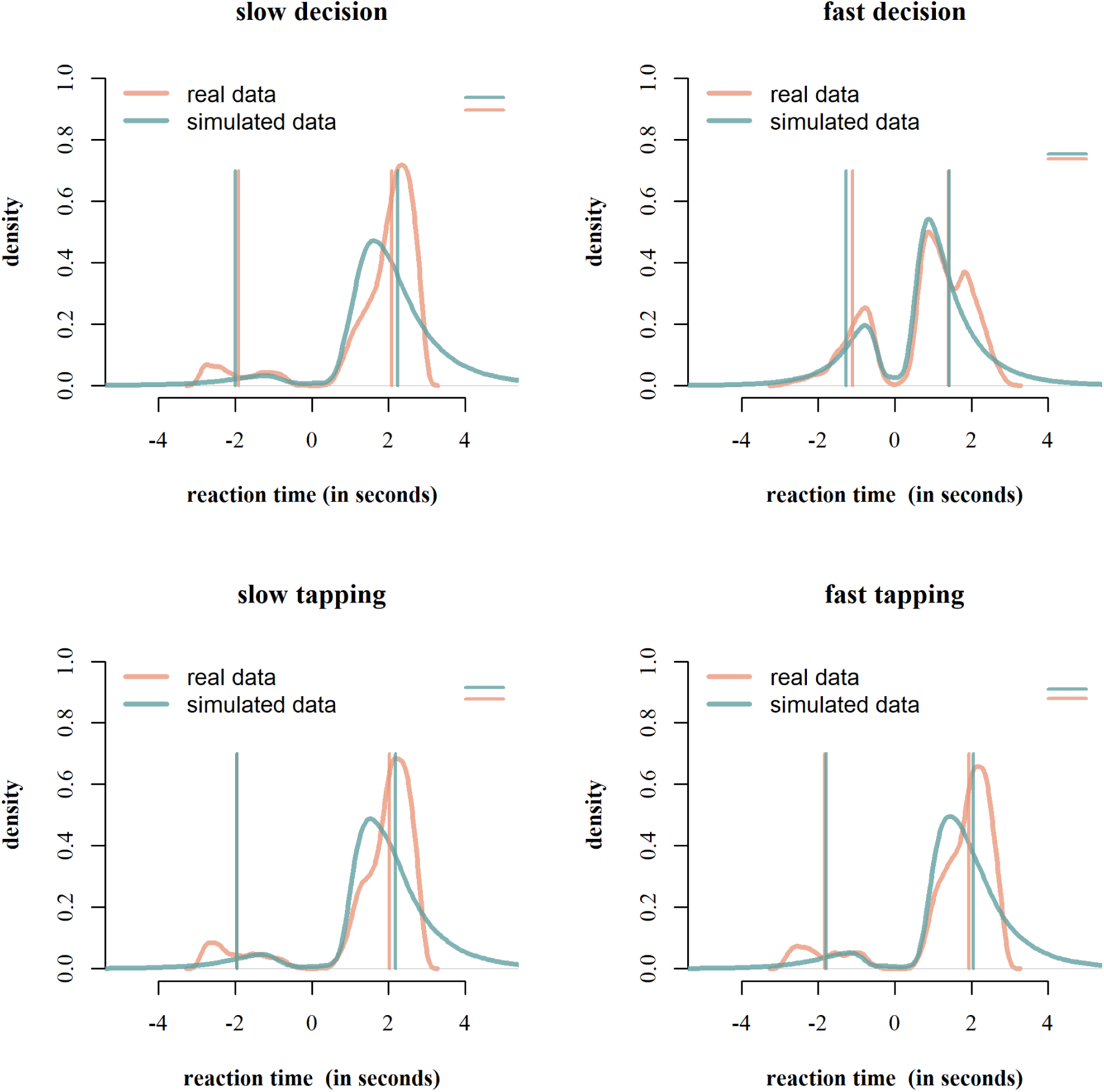
Comparison between empirical data and data as predicted by the best-fitting drift diffusion models of each session. Density distributions are plotted with a fixed bandwidth of 100 ms. Negative reaction times reflect incorrect decisions. Fitted models predict mean reaction times in the decision phase of the task (vertical bars) and proportion of correct decisions (horizontal bars in the upper right corner of each plot, scale from 0 and 1) of each experimental condition well, despite of a slight tendency to overestimate accuracy. Importantly, fitted models predict a reduction in reaction times for both correct and incorrect decisions in both fast-decision and fast-tapping blocks as compared to slow-decision and slow-tapping blocks, as observed in empirical data. As such, behavioral differences in reaction times observed in the decision phases of experimental conditions were replicated well, which was the effect of interest. Therefore, agreement between observed and simulated data was considered satisfactory for the purposes of this study. However, whereas early responses show a strong agreement between simulated and real data, late responses do not: In this experiment, by design it was advantageous to respond shortly before the timeout of three seconds to maximize success probability, causing shifted reaction time distributions towards late responses in real data. The drift diffusion model, ‘blind’ to this deadline, simulates a distribution with late responses exceeding three seconds, thereby underestimating the high proportion of responses just before timeout. Remarkably, data in fast-decision blocks pose the exception, as here real and simulated data show a strong overlap. This is because only in fast-decision blocks choosing as soon as possible was the best strategy, leading to a low proportion of responses just before timeout. For each participant, the same number of trials were simulated as present in the real data. Data was simulated in this way 100 times, as if simulated participants performed the experiment 100 times. Resulting response distributions were averaged to produce most likely behavioral distributions for each model.

**Table S1.**
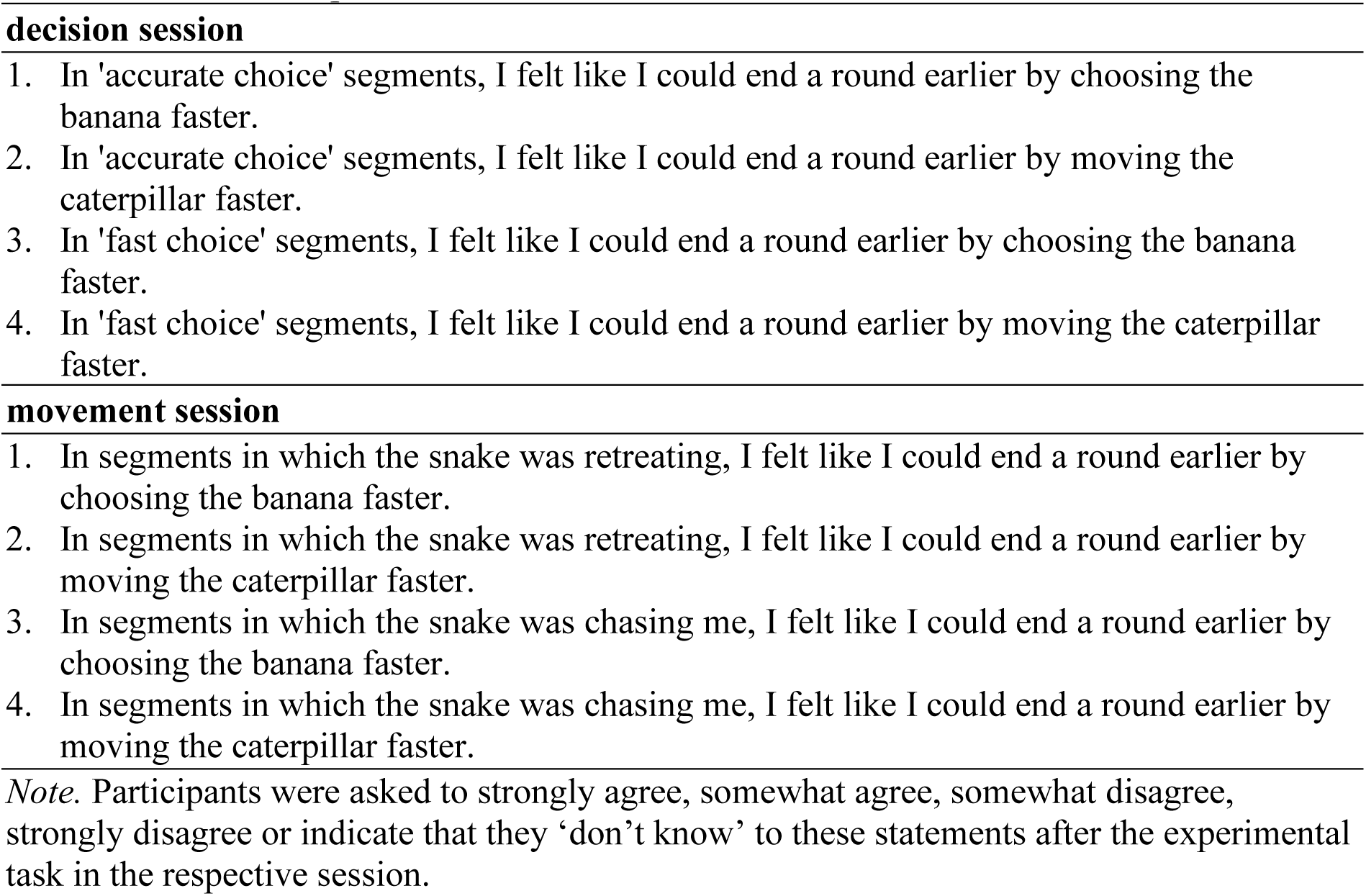
Instruction questionnaire.

**Table S2.**
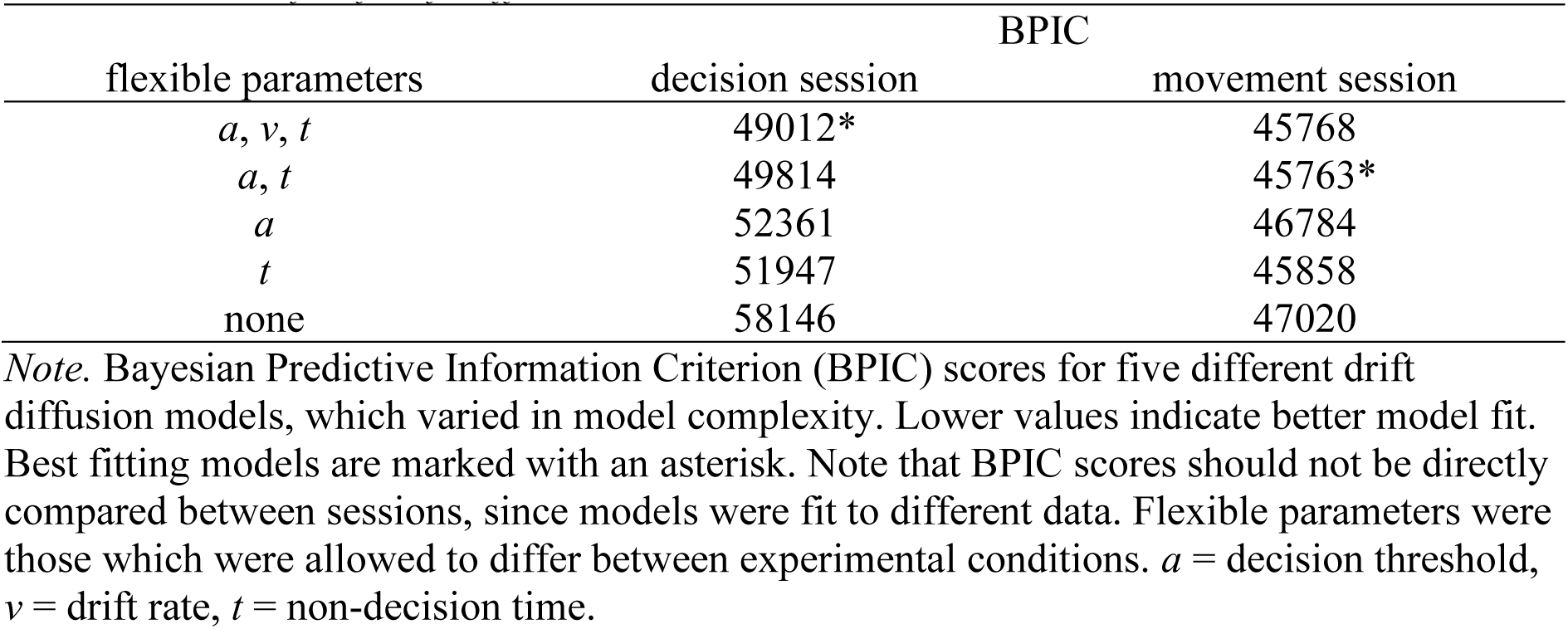
Model fit of drift diffusion models.

| flexible parameters | BPIC |  |
| --- | --- | --- |
|  | decision session | movement session |
| $a, v, t$ | 49012* | 45768 |
| $a, t$ | 49814 | 45763* |
| $a$ | 52361 | 46784 |
| $t$ | 51947 | 45858 |
| none | 58146 | 47020 |

*Note.* Bayesian Predictive Information Criterion (BPIC) scores for five different drift diffusion models, which varied in model complexity. Lower values indicate better model fit. Best fitting models are marked with an asterisk. Note that BPIC scores should not be directly compared between sessions, since models were fit to different data. Flexible parameters were those which were allowed to differ between experimental conditions. $a$ = decision threshold, $v$ = drift rate, $t$ = non-decision time.

